# Discovery of new marine species *Stentor hondawara* and its whole-genome reveal their unique ecology in comparison with freshwater stentors

**DOI:** 10.64898/2026.03.12.711380

**Authors:** Takato Honda, Daniel B. Cortes

## Abstract

*Stentor* is a genus of large ciliates that can be found in ponds, lakes, rivers, and fresh waters all over the world. Since their initial discovery in 1744, *Stentor* strains have been isolated from all populated continents. To date, over 50 individual strains have been identified, yet not a single isolate from a marine environment has been verified. Over 200 years since the initial description of the *Stentor* genus, our study entails the first concrete discovery of a fully marine *Stentor* species, as evidenced by its morphological, ecological, and phylogenetic positioning amongst *Stentor*. This new marine organism, which we have named *Stentor hondawara*, was verified to be a new species of the *Stentor* genus that appears to have fully adapted to a uniquely marine lifestyle in a high-salinity environment. Using comparative genomics analysis between the whole-genome sequences of *Stentor hondawara* and two freshwater species of *Stentor*, we further detected several intriguing differences in the enrichment of gene orthologs between the marine *Stentor hondawara* and the freshwater species, *Stentor coeruleus* and *Stentor pyriformis*. The gene groups specifically enriched in *Stentor hondawara* encode a variety of proteins, including ion channels, pH-responsive proteins, osmoprotectants, amino acid biosynthesis enzymes, and signaling receptors. Additionally, using metagenomics, we detected and isolated, from within our initial genome assembly, the genome of a novel marine bacteria, which we propose is an endosymbiont of *Stentor hondawara*. This bacterial species is an uncharacterized member of the order *Rhodospirillales* and appears to be a nutritional factory for the host *Stentor hondawara.* Taken together, our study provides insight into how *Stentor hondawara* adapted to a marine environment distinct from the habitats of all the other currently known *Stentor* species living in freshwater.

## Introduction

*Stentor* is an understudied ciliate genus characterized by large trumpet-shaped ciliated unicellular organisms capable of remarkable feats of regeneration and behavioral complexity. Like many other ciliate genera, *Stentor* can be found in nearly every type of freshwater, from small ponds to large lakes. Members of the genus *Stentor* are profoundly diverse in their lifestyles and can fill all manner of ecological niches, with different species blooming seemingly year-round. Since the genus was first described in 1744^1^, over fifty different strains have been isolated from all around the world^2^. Original morphological and behavioral characterization suggested a phylogeny of approximately fifteen to twenty unique species^2^, with several conspecific strains (divergent strains of the same species with some morphological variability) or species complexes (closely related taxa with highly similar morphologies, which may appear indistinguishable)^3^. More recently, use of the 18S SSU molecular marker, the evolutionary history of over 20 *Stentor* strains, has generated a phylogeny of approximately 12 true distinct species^4^, including the recently identified *S. stipatus*^5^. Only within the last 10 years have the genomes of any *Stentor* species been assembled, and only in the last two years have multiple partially annotated genomes been available for comprehensive genomic comparison^6,7^. Therefore, despite significant advancements in the last few decades, many questions remain about the evolutionary history, ecology, and biogeographic distribution of *Stentor*.

Like many other ciliate genera, *Stentor* is believed to be either exclusively or predominantly freshwater-dwelling^2,8^. Few mentions to marine “stentors” exist, but upon further study, none have been validated, with most being placed in morphologically-similar genera like *Condylostentor*^9^ or unverified^10^. Most recently, in 1996, large reddish ciliates were isolated from the reefs around coastal Guam. Upon initial inspection, the morphological characteristics of these ciliates suggested a close evolutionary relationship with *Stentor*. However, 18S SSU sequence analysis demonstrated that this organism was not a *Stentor,* but rather a new genus called *Maristentor*^11^. Thus, current consensus dictates that *Stentor* is exclusively a freshwater ciliate. Interestingly, recent efforts at sequencing environmental DNA (eDNA) from marine samples across the world have demonstrated a significant number of hits for *Stentoridae*, a larger taxonomy which includes *Stentor* and *Condylostentor*, a rare genus of marine ciliates, and a few potential hits to actual *Stentor* species (www.obis.org).

In this work, we introduce a novel *Stentor* species, *Stentor hondawara*, named for a group of marine brown kelp (known as ‘Hondawara’ seaweed in Japanese) upon which these cells were first isolated, and honoring their evolutionary relationship to its nearest known relative *Stentor katashimai,* a freshwater species found in Japan^12,13^. *Stentor hondawara* has been isolated over three consecutive summers from the coastal Atlantic Ocean waters of Cape Cod in Massachusetts. Here, we provide an initial morphological description of *Stentor hondawara*, along with a phylogenetic comparison to other *Stentor* species using the 18S SSU molecular marker. We include the whole genome, assembled from next-generation sequencing with short paired reads, and protein predictions generated using the partially annotated genomes of *S. coeruleus*^6^ and *S. pyriformis*^7^ as references for GeMoMa^14,15^ prediction. Finally, we perform comparative genomics through functional annotation and orthogroup enrichment comparison between *Stentor hondawara* and the two freshwater species to provide insights into some key differences between freshwater and marine *Stentor* species.

## Results

### Stentor hondawara taxonomy

The ZooBank registration number of this work is: urn:lsid:zoobank.org:pub:CCA05001-C7DD-46E5-8E07-BD0837CD9AB2.

The species described in this work, *Stentor hondawara*, is registered on ZooBank: urn:lsid:zoobank.org:act:B030D29A-470F-4DFC-A6A6-5AC647E373E6.

#### Diagnosis

Fully marine ciliate species with blue-green cortical pigmentation, often including reddish-brown patches (Fig. 1d). The macronucleus appears to be moniliform with approximately 6-10 nodes, though rich pigmentation makes it difficult to count all nodes (Fig. 1e-h). Cells are approximately 500-700 microns long and 200-300 microns wide when fully extended (Fig. 1e-i; n = 6 individual live cells). Cells are often bell-shaped when attached to a substrate (Fig. 1f, i). Upon contraction, the membranelles retract fully (Fig. 1g, j-l), similar to what has been reported for *Stentor introversus.* 18S SSU sequence comparison places this organism amongst *Stentor*. The pigmentation, size, and adaptation to a fully marine environment all suggest this marine *Stentor* is a distinct species.

**Figure 1.**
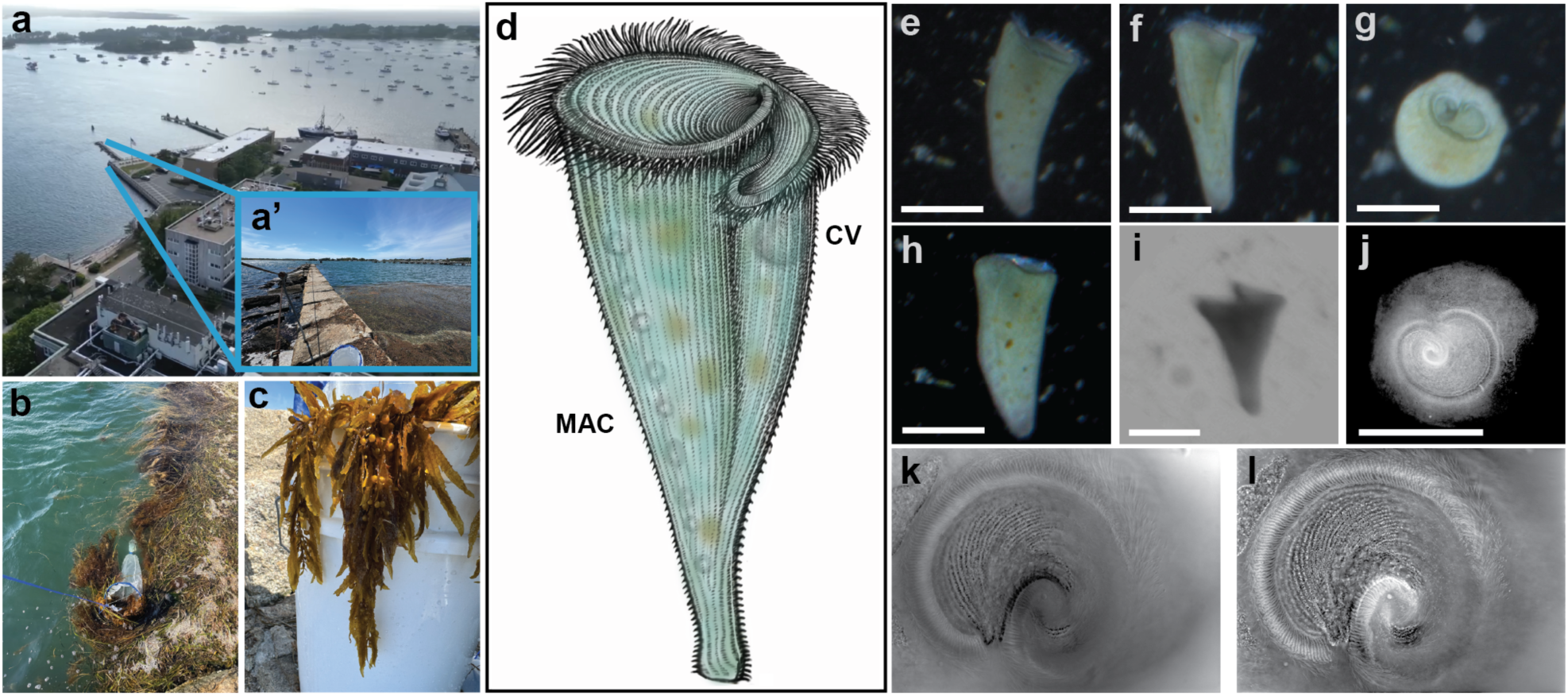
Collection and morphology of *Stentor hondawara.* **a,** Aerial overview of the Marine Biological Laboratory (MBL) campus in Woods Hole, MA. Blue inset **a’** is the jetty behind the National Oceanic and Atmospheric Administration (NOAA) parking lot where samples of *Stentor hondawara* were collected (Supplementary Information Video S1). **b,** Photo of the collection process, whereby a plankton net is towed 1-10 feet below the surface all along the jetty. **c,** Representative sample of *Sargassum* collected along with water samples. **d,** Detailed colored drawing of *Stentor hondawara*. The contractile vacuole (CV) and macronuclear nodes (MAC), which are sometimes visible by darkfield imaging, are shown in their relative positions in the sketch. The body is a green-blue color with brown-red stain spots near and on the surface, possibly food vacuoles and or the cultures of prospective endosymbiont bacteria. **e–f,** Darkfield image of a *Stentor hondawara* cell in swimming form taken from timelapse videos (Supplementary Information Video S2-S4). **g,** Darkfield image of a *Stentor hondawara* cell contracted, taken from a timelapse video. **h,** The same cell from e-f a few frames later, where the MAC nodes are more visible along the left side of the cell, and the membranelles are fully retracted. **i,** Transmitted light image of a different *Stentor hondawara* cell in attached form, with visible sharp cleft separating the bi-lobed frontal field. **j,** Transmitted light image of the frontal field of a contracted *Stentor hondawara* cell, showing retracted membranelles. **k,** OI-DIC image of the frontal field of a fixed contracted *Stentor hondawara* cell, showing the adoral membranelles spiraling towards the cytostome opening of the cell. **l,** OI-DIC image of the same cell as in (**k**) but at a different focal plane, highlighting the membranellar bands and pigment granules that fill the spaces between the adoral membranelles. All scale bars are approximately 250 microns.

#### Type specimens and locality

DNA was isolated from ∼300 sub-clonal cells and used to assemble the whole genome, which is deposited in the NCBI database (submitted and awaiting Accession ID). The original gDNA samples of *Stentor hondawara* have been secured, stored, and maintained in a -80 °C deep freezer at MIT (Cambridge, MA 02139, USA), which is available from the corresponding authors upon reasonable requests. For morphological analysis, slides were prepared with either cells fixed in glutaraldehyde or live cells suspended in filtered Pacific Ocean Water (fPOW; Imaginarium). The hapantotype, consisting of a slide with glutaraldehyde-fixed cells, has been stored in a 4 °C refrigerator at the Cortes lab at Virginia Tech (Blacksburg, VA 24061, USA).

#### Ecology

This new species was found in marine samples only on sunny days when the ocean water temperature ranged between 20-22 °C. The ocean salinity in the collection area was found to range between 35-40 ppt. The best predictors of *Stentor hondawara* presence appeared to be coincident with a heightened presence of brown kelp *Sargassum* and ctenophores *Mnemiopsis leidyi. Stentor hondawara* cells have been found to anchor on the kelp (Fig. 1c) as well as on the shells of small marine snails (species unknown). In culture, *Stentor hondawara* cells are voracious eaters and grow readily in 60 mm petri dishes if given fresh ocean water directly from the MBL coastline every 2-3 days. Although we collected ∼800 live *Stentor hondawara* individuals in this study, attempts to grow this species away from the MBL have thus far been thwarted by an inability to establish optimal culturing methods.

#### Etymology

The species name, *Stentor hondawara*, is based on the Japanese noun “Hondawara”, meaning the seaweed group *Sargassaceae* (family) and *Sargassum* (genus), where *Stentor hondawara*’s major habitats are in natural seawater (Fig. 1c). The authors recognized and utilized the Hondawara marine algae as a key landmark for specimen collection. Hondawara is known as a lucky charm for a good harvest in Japan, meaning ‘rice in the sea’ because of its similar shape along with air bladders (Fig. 1c), and it functions as a vital habitat for various marine species. Japanese name: Hondawara-rappa-mushi.

#### Morphology

*Stentor hondawara* cells were easy to discern from surrounding microorganisms by their trumpet-shaped, corkscrew swimming pattern and blue-green pigmentation (Fig. 1d-h). Under dissection stereomicroscope live imaging, *Stentor hondawara* cells generally appeared to have uniform pigmentation with patches of reddish-brown color, and lateral striations just barely visible (Fig. 1d), very similar to *S. coeruleus*. Under darkfield imaging, however, pigmentation became a lighter teal color (Fig. 1d-h), unlike *S. coeruleus,* whose pigmentation can often look purplish or dark blue under darkfield imaging. When fully extended, *Stentor hondawara* measured approximately 500-700 microns in length by 200-300 microns in width (n = 6 live cells). Imaging of glutaraldehyde-fixed cells by OI-DIC^16^ revealed approximately 25-30 peristomial kineties (n = 3) and 210-260 adoral membranelles, or 7-8 per 10 microns (n = 3), with the average oral apparatus being approximately 250 microns wide (n = 4). The macronucleus (MAC), which was visible under oblique lighting conditions in live cells and in some fixed cells (Fig. 1d and 1h), appeared to have a moniliform organization with around 6-10 total nodes, though it remained difficult to count the complete number of nodes due to the heavy pigmentation of the cell. When attached to a substrate, *Stentor hondawara* cells often adopted a sharper, more pyramidal shape, retaining their extended oral apparatus and frontal field. When contracted (Fig. 1g), *Stentor hondawara* cells became almost spherical (∼400 microns long by ∼300 microns wide) (n = 5 live cells and n = 4 fixed cells) and seemed to retract their membranelles internally (Supplementary Information Video S3). A partial retraction of the membranelles was also seen to occur periodically in attached cells while feeding normally (Supplementary Information Video S2, S4).

### Stentor hondawara phylogeny

The *Stentor hondawara* 18S SSU sequence (2,044 bp) was found by BLASTn search against the full *S. pyriformis* SSU sequence. Using 18S alignment with *Maristentor dinoferus*, *Blepharisma americanum,* and *Condylostentor auriculatus* as outgroups, we estimated a phylogenetic topology that clustered *Stentor hondawara* within *Stentor.* Furthermore, *Stentor hondawara* appeared to share a common ancestor with *S. katashimai*, *S. polymorphus*, *S. igneus*, *S. muelleri*, and *S. roeselii* (Fig. 2). The tree topology suggests *S. katashimai* as the closest relative to *Stentor hondawara*. Based on UfBoot, aBayes, and SH-aLRT (75.8/0.968/80), along with the overall branch length between the two samples, and its unique ecology, *Stentor hondawara* is a novel species. Importantly, our tree topology maintains established relationships between different *Stentor* clusters, such as the clustering of *S. roeselii* and *S. muelleri* into a single node, and the clustering of *S. stipatus, S. amethystinus,* and *S. pyriformis* into a separate node (Fig. 2). Thus, *Stentor hondawara* is a novel species of *Stentor* that has entirely adapted to marine life, unique amongst known *Stentor* species.

**Figure 2.**
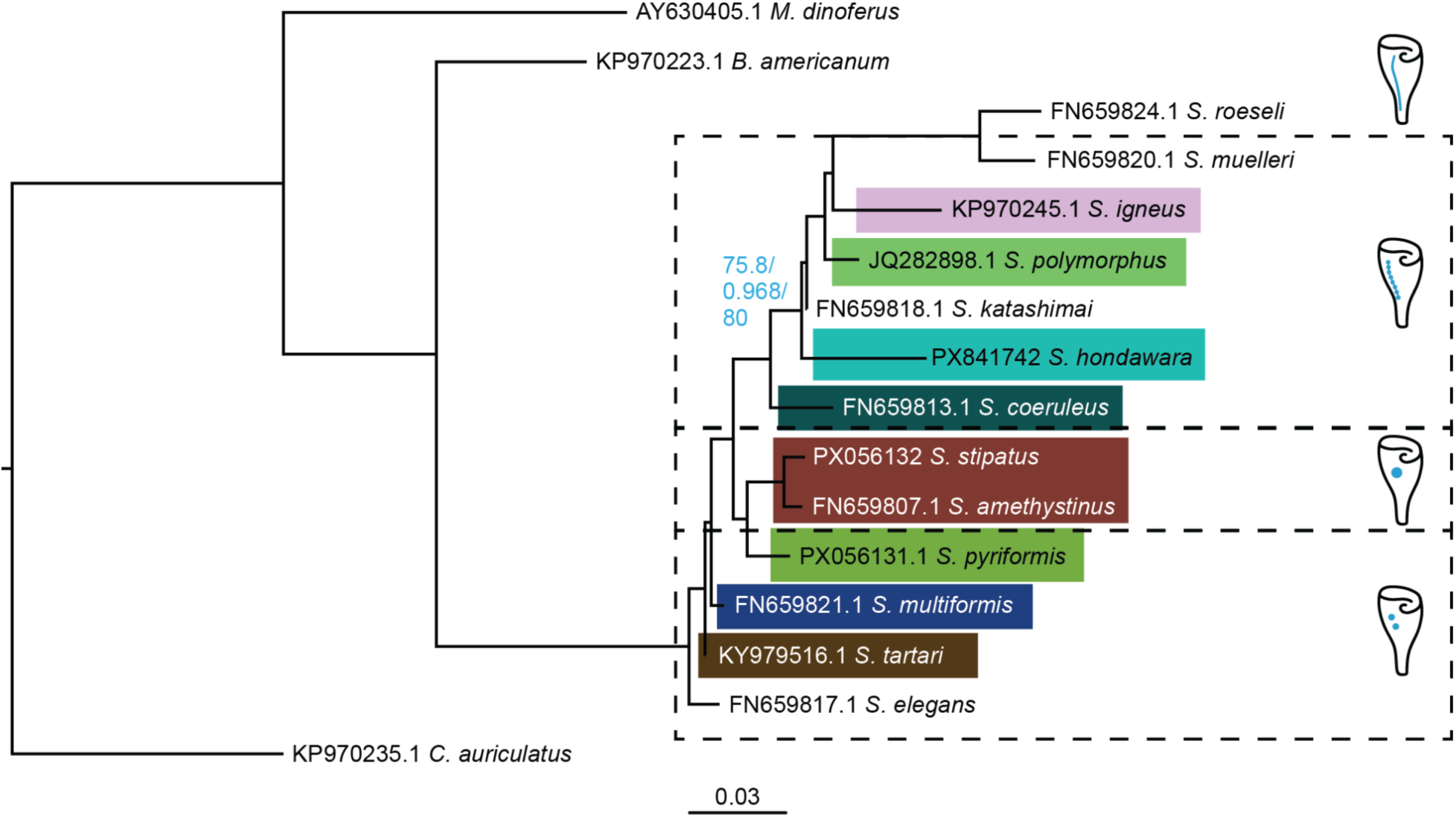
*Stentor hondawara* in the *Stentor* phylogeny. Phylogenetic tree assembled using maximum likelihood (ML) estimates, with branch lengths showing nucleotide substitutions per site. Blue text highlights the reliability measure scores (UfBoot/aBayes/SH-aLRT) for the branch separating *Stentor hondawara* from *Stentor katashimai* and the rest of that node. Colored boxes highlight the pigmentation or otherwise coloration of each *Stentor* species. Cartoons on the far right denote the general macronuclear structure associated with each sub-group of *Stentor* species, separated by dashed lines. The scale bar indicates nucleotide substitutions per site.

### The *Stentor hondawara* macronuclear genome

We tried both *de novo* (SPAdes and MEGAHIT)^17,18^ and referenced (MaSuRCA)^19^ genome assembly with *S. coeruleus* as the reference genome. However, the referenced assembly failed to yield an assembly (see also ‘Methods: Whole-genome sequencing and raw assembly of *Stentor hondawara* genomic DNA’). In contrast, *de novo* assembly yielded high quality complete genomes, with SPAdes outperforming MEGAHIT by N50 and contig number. The raw assembled *Stentor hondawara* macronuclear genome was 98.3 Mbp long with an N50 of 55.1 kbp, consisting of 183,448 total contigs. BlobTools2 was used to filter out contaminants from the genome assembly by size, coverage, and GC% (Fig. 3 ‘*Stentor hondawara* genome’ cluster and Extended Data Fig. 1). The remaining contigs included one very large *Ciliophora* contig with GC content around 13% and coverage over 1000x, and several large *Pseudomonadota* contigs with GC content around 45% and coverage around 200-300x. Both of these clusters were filtered out and separated, the first as the putative mitochondrial genome, and the second as a putative endosymbiotic bacterial genome.

**Figure 3.**
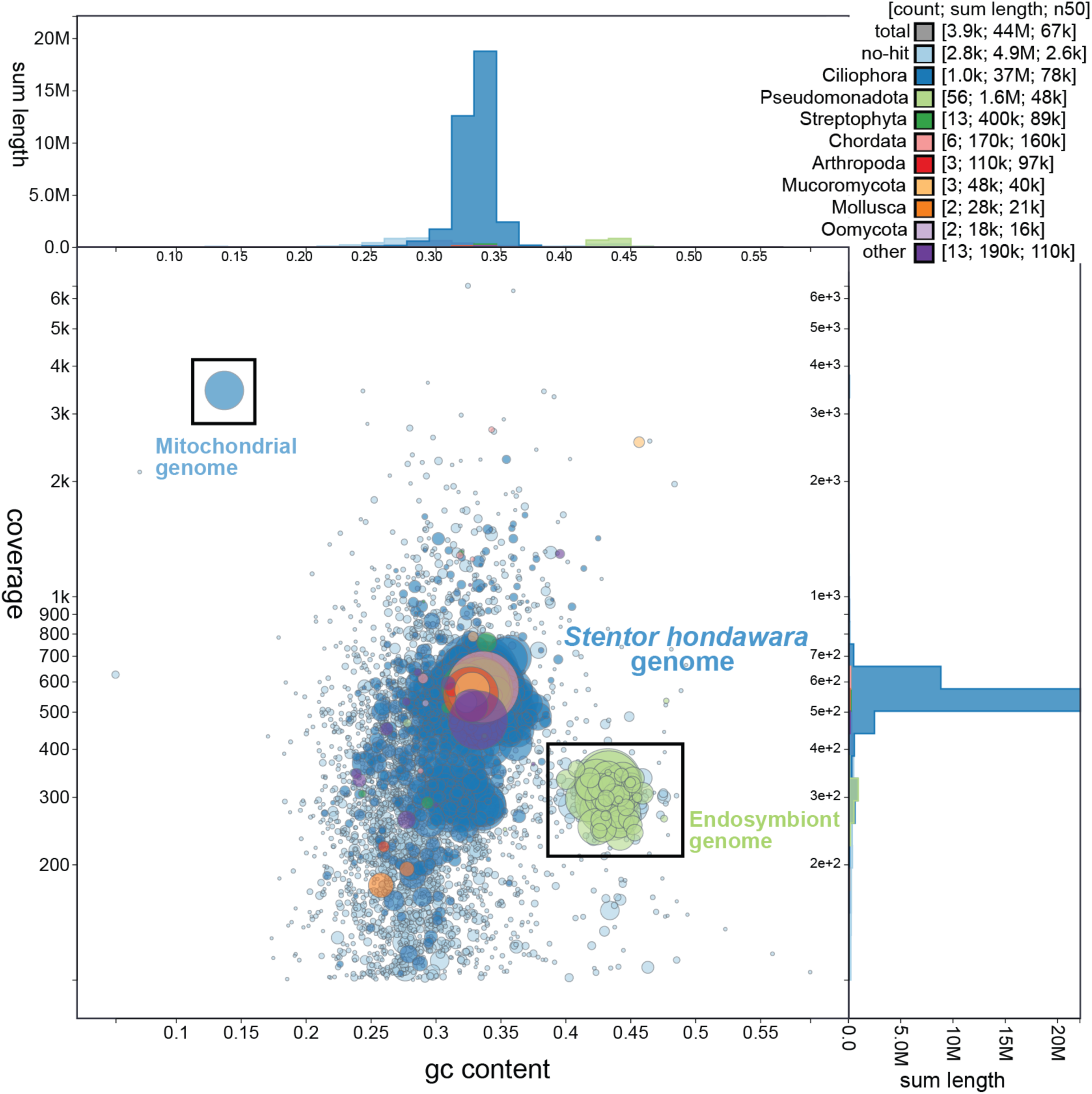
Analysis of the *Stentor hondawara* hologenome. Distribution of all contigs found in the partially filtered *Stentor hondawara* genome, where contigs with less than 100x coverage and smaller than 200 bp in length have been removed. Each circle denotes a single contig whose color specifies the highest score taxonomic hit for that contig. Note the single large contig with extremely low GC content, which we identified as the entire mitochondrial genome, and the cluster of *Pseudomonadota* and “no-hit” contigs with significantly higher GC content than the main genome, which we identified as a potential endosymbiont. This bacterial genome was filtered out from the final decontaminated assembly. The plot was made with Blobtools2.

The resultant filtered *Stentor hondawara* genome consisted of 3,706 contigs with a total genome length of 41.7 Mbp, an N50 of 68.9 kbp, GC content of 32.66%, and 182.7 N’s per 100 kpb on average (Extended Data Fig. 1-2). This assembly was found to contain 97.7% complete BUSCO orthologs [with 78.4% single-copy and 19.3% duplicates], 1.8% fragmented BUSCO orthologs, and 0.5% missing orthologs compared to the alveolata_odb10 dataset. These BUSCO scores were consistent with the scores achieved by the published *Stentor coeruleus* genome (96.5% complete *Alveolata* BUSCOs), suggesting a similar level of completeness amongst these genome assemblies. The final *Stentor hondawara* genome assembly is intermediate in size compared to the much larger (∼62.4 Mbp) *S. coeruleus* genome^6^ and the much smaller (∼29.2 Mbp) *S. pyriformis* genome^7^. Despite such differences in macronuclear genome size, mapping of raw *Stentor hondawara* reads to either *S. coeruleus* or *S. pyriformis* genomes using minimap2 and Bowtie2 demonstrated a higher percentage of total aligned reads to the *S. coeruleus* genome (4% vs 0.6%), suggesting higher similarity between the two.

The mitochondrial genome assembly, NODE_319 (contig ID), was found to be extremely AT-rich and 48,414 bp long (Fig.3 ‘Mitochondrial genome’ cluster). We verified that NODE_319 contained mitochondrial sequences by annotation with MFannot^20^ and Barrnap^21^. 42 total genes were identified, including 27 protein-coding genes: atp9, cox1, cox2, cob, nad1, nad4, nad4L, nad5, nad7, nad9, nad10, several rpl proteins, several rps proteins, and several unknown hypothetical proteins (Extended Data Table. 3). Barrnap also discovered two ribosomal RNA genes, rnl and rns, and 5 tRNA genes (Extended Data Table. 3). Several genes were missing, including cox3 and most tRNA genes, though we note that nearly all of the same genes (including cox3) appear to be missing from the published complete mitochondrial genome of *S. coeruleus.* We therefore concluded that this single large contig contains the majority, if not the entirety, of the *Stentor hondawara* mitochondrial genome.

### Annotation of the *Stentor hondawara* genome

We predicted the protein-coding genes present in the *Stentor hondawara* genome using GeMoMa^14,15,22^, using both *S. pyriformis*^7^ and *S. coeruleus*^6^ gene annotations as references for gene prediction. This resulted in an annotation of 20,565 total predicted protein-coding genes. Analysis revealed a total of 5,873 introns mapped to these 20,565 protein-coding sequences, for an average of 0.29 introns per gene. Looking more closely, of the total protein-coding sequences, only 4,373 contained introns (21.3% of all predicted protein-coding genes) and averaged 1.34 introns per gene. Most introns found were around 30-50 bp with a median intron size of 45 bp. Using eggNOG-mapper2^23^ with the eggNOGv5.0 database^24^, we next looked for functional annotations by DIAMOND^25^ protein sequence homology with the BLOSUM45 protein alignment matrix. This resulted in the functional annotation of 4,462 genes, or approximately 21.7% of all predicted protein-coding genes.

### *Stentor hondawara* unique orthogroups

Given that all other known *Stentor* species are exclusively freshwater-dwelling, we were interested in finding any specific marine adaptations that could be inferred from comparative genome analysis between the *Stentor hondawara* genome and those of *S. coeruleus* and *S. pyriformis.* OrthoFinder^26^ analysis, run on all three sets of predicted protein-coding genes, revealed a set of 590 orthogroups that were unique to *Stentor hondawara* among the three *Stentor* species, with 9,787 orthogroups common to all three and 1,367 orthogroups unique to both freshwater species (Fig. 3a). eggNOG^22^ analysis of the 590 *Stentor hondawara-*specific orthogroups revealed hundreds of significantly-enriched gene groups with predicted functions including GPCRs, response to light stimulus, response to low-fluence blue light, organic/amino acid: sodium symporters, cellular response to pH, cellular response to potassium ion starvation, transmembrane transporters, carboxylic acid transmembrane transporters, voltage-gated chloride channels, chloride transporters, fatty acid homeostasis, fatty acid biosynthesis regulation, and fatty acid transport (Fig. 4a-b and Extended Data Table. 1). We compared these results to eggNOG analysis of the freshwater-specific orthogroups (Fig. 5a-b and Extended Data Table. 2) where we found enrichment of different orthogroups. Visualization of specific enriched groups in *S. coeruleus* vs *Stentor hondawara* demonstrates different ion regulation and signaling strategies (Fig. 4b). While the *S. coeruleus* genome encodes far more enriched sodium ion transport proteins, *Stentor hondawara* instead encodes more enriched potassium and chloride ion transport proteins. Similarly, *S. coeruleus* has a preference for cAMP enrichment while *Stentor hondawara* prefers MAPK and GPCR enrichment (Fig. 4b, 5a).

**Figure 4.**
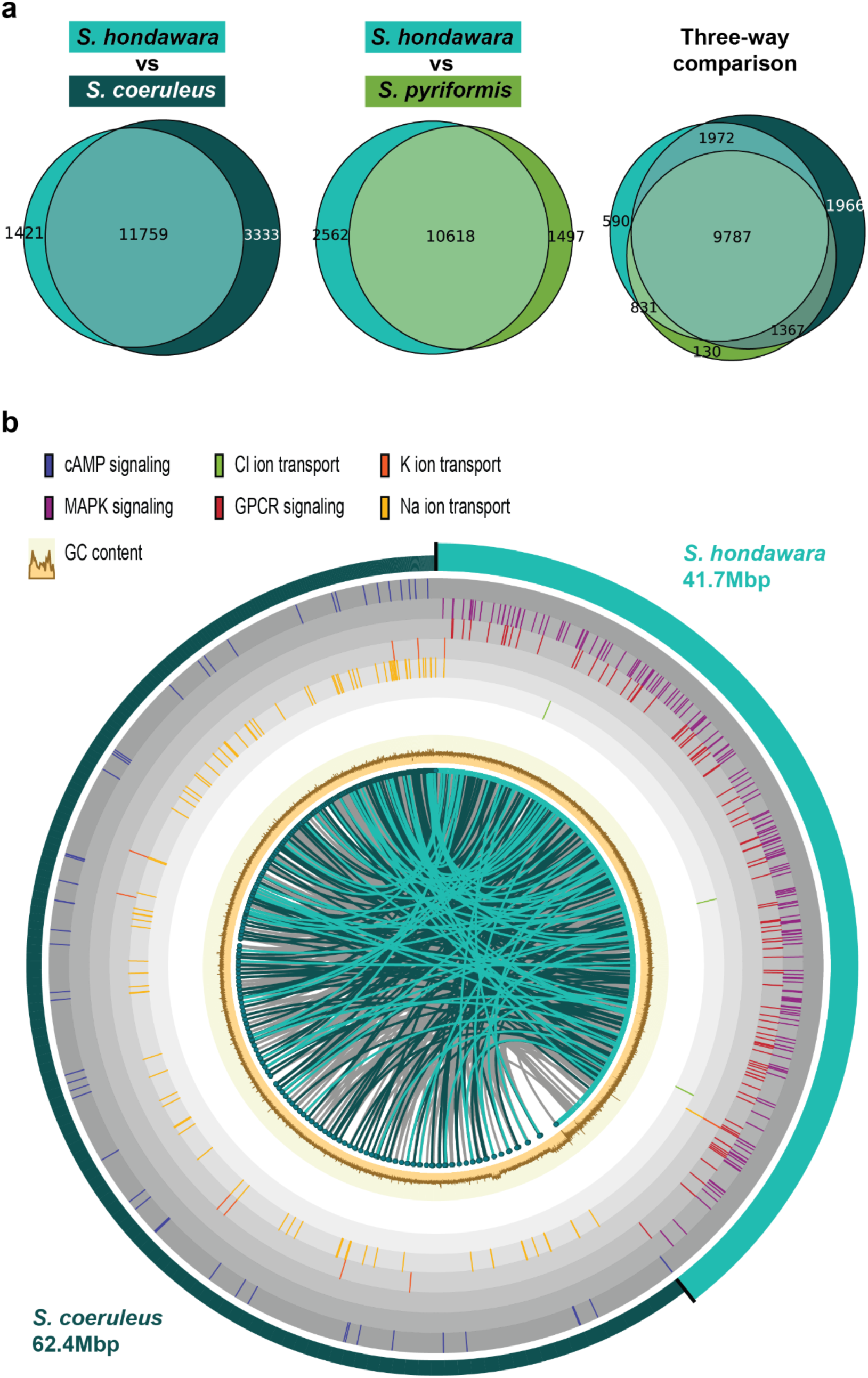
Ortholog comparison between *Stentor hondawara* and freshwater species. **a,** Venn diagrams showing pairwise comparison between the identified gene ortholog groups that are common or unique between *Stentor hondawara* and *S. coeruleus* (left) or between *Stentor hondawara* and *S. pyriformis* (middle), or between all three species (right). The size of the circles is proportional to the overall number of predicted genes in the structurally annotated genome of each species. **b,** A circular diagram representing the whole genome assemblies of *Stentor hondawara* and *S. coeruleus* (outermost solid-colored segments). Mapped genes (inner circle of colored dots) represent the gene orthologs that are common between both freshwater (*S. coeruleus* and *S. pyriformis*) and marine (*Stentor hondawara*) species. Lines linking these dots represent the links between the corresponding orthologs; grey-colored lines are generic genes that are not enriched in either freshwater or marine species. Lines colored to match either *Stentor* species color represent enrichment in that species over the other. The brown/tan plot just outside the gene orthologs represents the GC content estimated for both genome assemblies. Outer color-coded lines highlight the position of functionally-annotated GO (Gene Ontology) gene orthologs (color-coded by GO group), showing differential enrichment in *Stentor hondawara* versus *S. coeruleus*.

**Figure 5.**
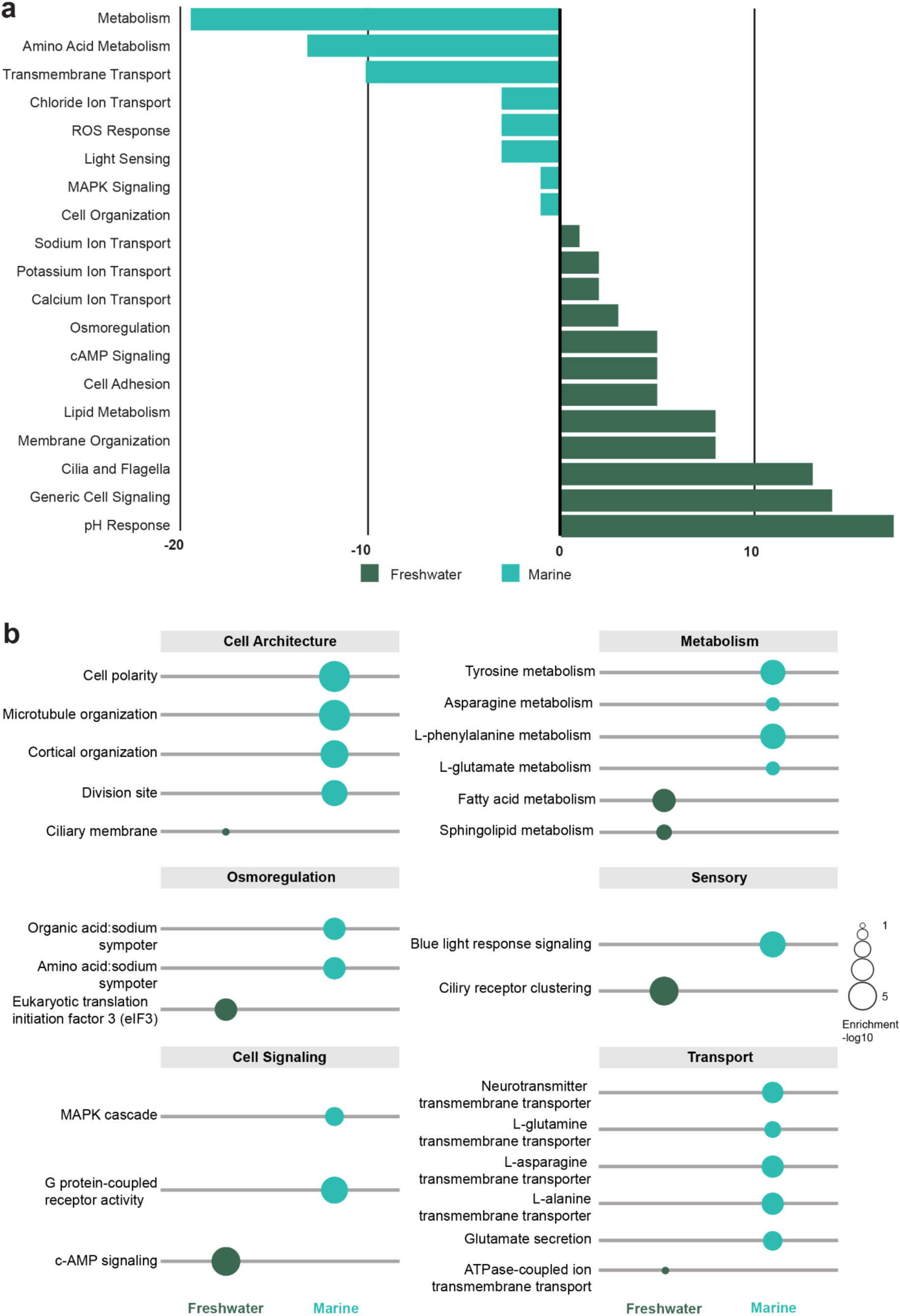
Marine and freshwater enriched orthogroups. **a,** Plot highlighting enriched functional orthogroups that are either freshwater-exclusive (positive, dark blue-green) or marine-exclusive (negative; light blue-green). Enrichment is shown as the total number of exclusively freshwater orthogenes minus the total number of exclusively marine orthogenes for each functional category. **b,** Detailed breakdown of gene groups enriched in several categories. The size of the representative circle for each pathway or process indicates enrichment over other lifestyles (freshwater or marine).

### Several predicted *Stentor hondawara* proteins suggest adaptations to marine life

Of the many gene orthologs enriched in *Stentor hondawara,* several had predicted functions that could assist in marine life adaptation. Among the many enriched ion channels predicted by eggNOG annotation, voltage-gated chloride ion channels of the CLCN2 family came up several times. In yeast, CLCN2 ortholog GEF-1 is a type of ion channel involved in cation regulation^27^, while in mammals, CLCN2 can be activated by hyperpolarization^28,29^, cell swelling^29,30^, extracellular hypotonicity^30^, and extracellular acidification^29,31^. We therefore identified the *Stentor hondawara* CLCN2 ortholog with the highest protein sequence identity to human CLCN2 by BLASTp search. Superimposition of human CLCN2 and the *Stentor hondawara* CLCN2-like protein predicted 3D structures in PyMol (Fig. 6b-b’; Supplementary Information Video S5) resulted in an alignment score of 1810.065 with an RMSD of 2.447. Interestingly, the *Stentor hondawara* chloride ion channel appears to have several longer alpha helices than the human ortholog.

**Figure 6.**
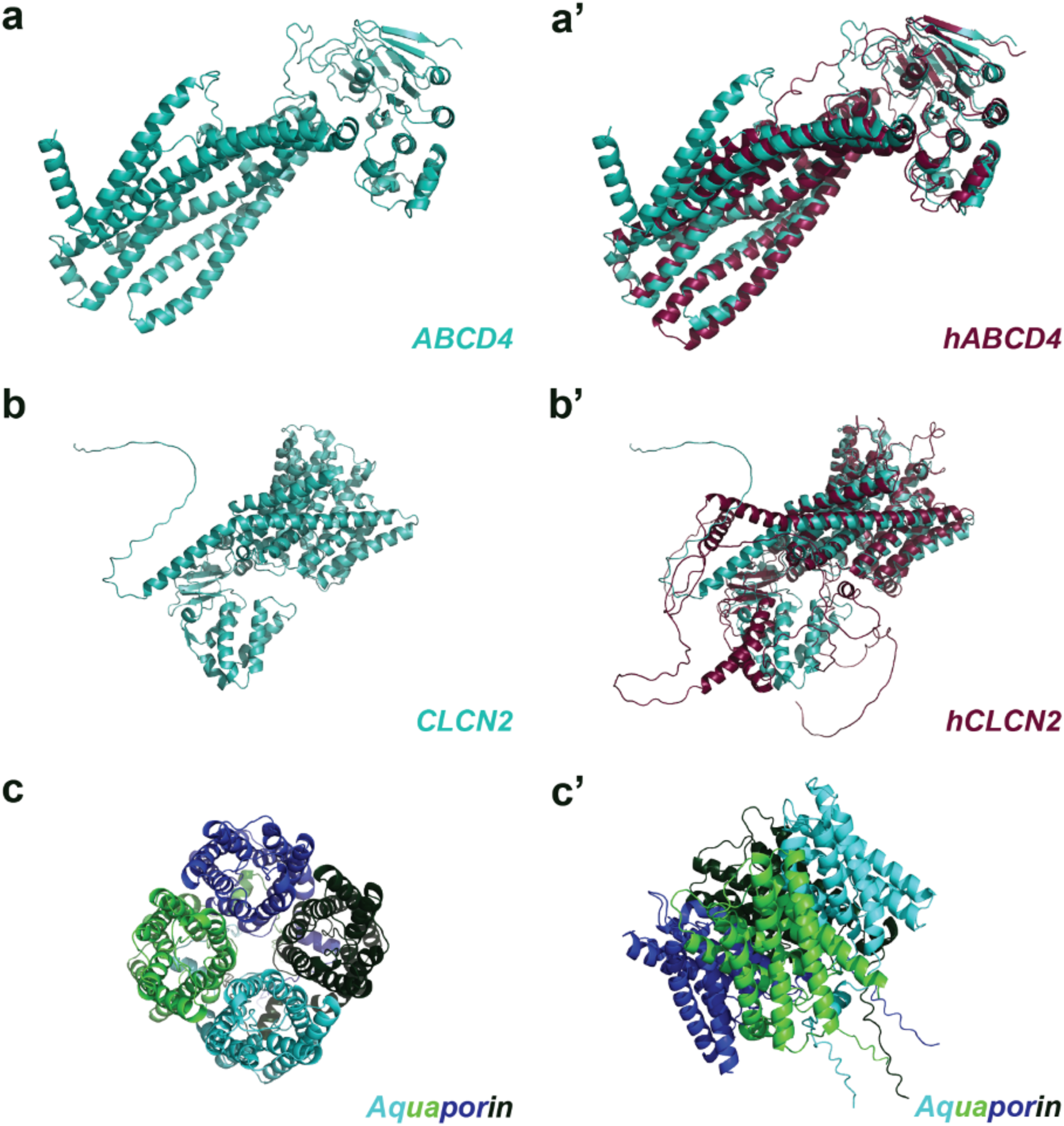
3D structures of uniquely identified proteins in *Stentor hondawara.* **a,** Alphafold-predicted structure of the top *Stentor hondawara* protein match for human ABCD4 (a vitamin B12 transporter) by protein sequence. **a’,** The alphafold-predicted structure of the human ABCD4 overlaid atop the *Stentor hondawara* ortholog. **b,** Alphafold-predicted structure of the top *Stentor hondawara* protein match for human CLCN2 (a voltage-gated chloride ion channel) by protein sequence. **b’,** The alphafold-predicted structure of the human CLCN2 protein overlaid atop the *Stentor hondawara* ortholog. **c,** Alphafold-predicted structure of a *Stentor hondawara* ortholog to human Aquaporin-7, an aquaglyceroporin, showing the predicted homotetrameric structure of the complex. **c’,** A different angle of the same predicted protein structure shown in (**c**). See also Supplementary Information Video S5-S7.

Given the general enrichment of osmoprotectant mechanisms in *Stentor hondawara*, we next looked for *Stentor hondawara* aquaporins, specifically aquaporin-3, as this is an important aquaglyceroporin that is important for osmoregulation in many marine organisms^32,33^. BLASTp search against human aquaporin-3 resulted in the identification of 13 separate *Stentor hondawara* aquaporins. We were interested in identifying whether any of these hits were specifically aquaporin-3-like, so we performed multiple sequence alignment of all aquaporin protein sequences using MAFFT^34^ (BLOSUM45 substitution matrix). In order to increase the resolution of the aquaporin phylogenetic tree and better discern aquaporin isoforms, we included all identified aquaporins for *Stentor pyriformis* (11 hits) and *Stentor coeruleus* (15 hits), as well as human, nematode, algal, yeast, and plant aquaporins. Interestingly, most *Stentor* aquaporins were represented in all three species, but two clusters were exclusive to *Stentor coeruleus,* and one was exclusive to *Stentor pyriformis* and *Stentor hondawara*. This *Stentor hondawara* cluster appeared to show two separate duplications of the gene, resulting in four total paralogs (Extended Data Fig. 3). The phylogenetic tree calculated from alignment of all mentioned aquaporins revealed that all *Stentor* aquaporins form a node separate from those of the more modern eukaryotic forms of life. We therefore used ProtSpace^35^, a protein embedder that reduces multidimensional data to cluster proteins of potentially similar function, to analyze whether any *Stentor* aquaporins clustered with aquaglyceroporins by dimensional reduction (t-SNE) (Extended Data Fig. 4). Indeed, when considering both local and global structure, the uniquely-enriched *Stentor hondawara* aquaporin paralogs were revealed to cluster with human and other eukaryotic aquaglyceroporins (Extended Data Fig. 4). Alignment mapping of conserved regions (FAT/FST, NAS, and Froger residues)^36,37^ using ESPript v3.2 demonstrated that the presumptive *Stentor hondawara* aquaglyceroporins conserved most key residues, although they were missing P1 of the five Froger residues and their FAT/FST motif is a bit more divergent than those of typical aquaglyceroporins (Extended Data Fig. 5). Finally, we generated the predicted homotetramer structure of one of these predicted *Stentor hondawara* aquaglyceroporins, demonstrating a protein complex that closely resembles the known aquaporin complexes of other eukaryotes (Fig. 6c-c’; Supplementary Information Video S7). These data suggest that *Stentor hondawara* may have adapted a more generic aquaporin to function as a unique aquaglyceroporin, and then duplicated this adapted protein for better osmoregulation.

### Putative endosymbiotic bacteria in the *Stentor hondawara*

The *Pseudomonadota* cluster, along with all co-clustered “no-hit” contigs that contained few or no *Ciliophora* gene hits (< 50%), were filtered into a separate assembly totaling 249 contigs with a length of 2.3 Mbp (Extended Data Fig. 6 and Fig. 3 ‘Endosymbiont genome’ cluster). This assembly contained 2,058 predicted genes, including both 16S and 18S rDNA genes, based on Prokka^38^ annotation. BLASTx search demonstrated that most genes were closest to members of the order *Rhodospirillales* by sequence conservation. The bacterial CobN, CobT, CobS, and SecD protein sequences were isolated from the genome and concatenated to generate an estimated *Rhodospirillales* phylogeny (Supplementary Information Data S2). This phylogeny showed our putative endosymbiotic bacteria sharing a common ancestor with the families *Thalassobaculaceae* and *Aestuariispiraceae* (Extended Data Fig. 7). Nearly all known species in these two families appear to live in marine environments^39–42^. BUSCO analysis against the rhodospirillales_odb12 dataset demonstrated 87.1% complete [with 87.1% single-copy], 2.6% fragmented, and 10.3% missing BUSCOs. We therefore concluded that these 249 contigs denote a high-quality genome assembly (Extended Data Fig. 6) (2.3 Mbp; ∼200-400x coverage; N50 of 33 kbp; GC content of 43.37%, and 133.41 N’s per 100 kbp) of a potential endosymbiotic bacteria of the order *Rhodospirillales.* This bacterial genome has been uploaded to the NCBI database and is available through Accession ID (submitted and awaiting Accession ID).

Of the 2,058 total protein-coding genes predicted by Prokka^38^, 1,558 were assigned functional annotations (Extended Data Table. 4). As *Rhodospirillales* are potential nitrogen fixers^43–46^, we sought out nitrogen fixation genes (nifU, nifD, nifK, and nifH) within the predicted proteome. Hits were identified by BLASTp to nifU, nifK, and nifH, with several partial hits to nifD also present. Furthermore, we also identified several genes involved in Vitamin B12 (cobalamin) biosynthesis (Extended Data Table. 4; over 20 genes including cobQ, cobT, cobS, cobA, cobB, cobN, cobH, cobL, cobM, cobK, cobD, cobP, cobU, cobO, cobC)^47^, genes involved in respiration and energy production (including cytochrome C genes ctaC, ctaD, ctaB, ctaE, ctaG, ATP synthase, NADH dehydrogenase complex I, and nitrate reductase napA), amino acid biosynthesis (pathways for arginine, lysine, threonine, methionine, cysteine, histidine, tryptophan, phenylalanine, tyrosine, leucine, isoleucine, valine, proline, and serine), biosynthesis of several other key vitamins or cofactors (riboflavin, thiamine, pyridoxine, folate, NAD, heme, and ubiquinone), and carbon metabolism (glycolysis, gluconeogenesis, glyoxylate shunt, and TCA cycle genes). Several important genes for free-living *Rhodospirillales* appear to be missing from the annotation, including genes for biofilm formation, quorum sensing, and sugar transporters of the phosphotransferase system, along with fragmented or partially missing genes for flagella synthesis, and partially missing stress-response sigma factors (Extended Data Table. 4).

### *Stentor hondawara* may benefit from its endosymbiotic bacteria

Given the presence of an intact cobalamin biosynthesis pathway in the putative endosymbiotic *Rhodospirillales*, and since archaea and bacteria are the main marine source of this necessary nutrient^48^, we were interested in whether cobalamin transporters ABCD4 and LMBD1^49,50^ – two proteins important for transport of cobalamin into the cytosol, could be found in the *Stentor hondawara* genome. BLASTp search against both human orthologs yielded 27 hits for ABCD4 (Fig. 6a-a’; Supplementary Information Video S6) and two for LMBD1. *Alphafold3* predicted structures suggested strong structural conservation for both proteins. Superposition alignment in PyMol yielded an alignment score of 2107.337 with an RMSD of 2.57 for ABCD4 and an alignment score of 1366.759 with an RMSD of 2.851 for LMBD1. These alignment scores suggest that *Stentor hondawara* has functional transporters and is capable of utilizing cobalamin acquired from its endosymbiotic bacteria. We wanted to know whether either transporter was enriched in *Stentor hondawara* compared to freshwater *Stentor.* For ABCD4, we identified 27 hits in the *Stentor pyriformis* assembly and 41 in the *Stentor coeruleus* assembly (although the *S. coeruleus* genome has recently been suggested to have undergone whole genome duplication)^51^. Therefore, there is no significant enrichment for these importers in any *Stentor* – barring the enrichment by whole genome duplication. LMBD1, on the other hand, yielded only a single hit for either of the other two *Stentor* species, suggesting a slight enrichment in *Stentor hondawara*.

## Discussion

Here we introduce a novel species, *Stentor hondawara,* which is unique as the only currently established fully marine species of the genus. The few mentions that exist for any other potentially marine or partially marine species have been demonstrated to be either not *Stentor*^9,11^, or to be unverified, with no specific description or species diagnosis to back their claims^8^. We have preliminary evidence of another marine species of *Stentor*, more closely related to *S. muelleri* by 18S sequence alignment, which lives in brackish coastal waters of lower salinity (15-20 ppt compared to the usual 30-35 ppt for Atlantic Ocean water) and may not be fully marine. We are currently working on acquiring more samples of this other potential marine species so that we can sequence its genome and compare it to other known species, including *Stentor hondawara*.

Isolation efforts over three prior years have revealed that *Stentor hondawara* is a difficult species to find. Most freshwater *Stentor* species can be found at their source year-round, though they are sparse and therefore rare in samples most of the year unless they are actively blooming. In most cases, *Stentor* blooms occur between the early spring through late fall months. *Stentor hondawara*, on the other hand, is nearly impossible to find in the stated collection site for all but 1-2 weeks of each year – generally coinciding with water temperatures around 20-22 °C, the presence of large swaths of floating brown kelp of genus *Sargassum*, and significant presence of *Mnemiopsis leidyi* near the surface. *Sargassum* and *M. leidyi* species found along the North American east coast are generally migratory, usually moving up and down the east coast during the spring through fall months following temperate waters. *Sargassum* in the near coastal Atlantic settles into a region known as the Sargasso Sea^52^, which feeds the seasonal migrations from the Caribbean up through the northern east coast of North America and features vast microbial biodiversity^53^. As *Stentor hondawara* has not been isolated from the coastal shallows anywhere around Woods Hole, nor has it been isolated from the collection jetty or other nearby jetties aside from during the 1 to 2-week peak, it is very likely a migratory species. Initial collections of this *Stentor* species showed it primarily anchored along the stalks of *Sargassum* kelp or swimming primarily in water enriched for this kelp. It is therefore possible that *Stentor hondawara* makes its permanent home on this migratory kelp, feeding off the same microbial life that feeds the larval form of *M. leidyi*. Our initial attempt to test this idea, by trying to isolate *Stentor hondawara* from coastal marine samples along other regions of the North American east coast, resulted in the isolation of a different marine *Stentor* - but a single attempt at near-optimal temperature is not definitive and even at the MBL, not all days within the optimal temperature range resulted in detectable samples of *Stentor hondawara*.

The included assembly consists of the *Stentor hondawara* genome, its mitochondrial genome, and the genome of a novel *Rhodospirillales* bacterium, designated as a potential endosymbiont based on its genome coverage being similar to *Stentor hondawara* genome coverage but GC content being significantly higher. It is worth noting that while we are unsure if the *Stentor hondawara* mitochondrial genome is intact, it appears to be as complete as the published *S. coeruleus* mitochondrial genome, which is substantially larger, as both *Stentor* appear to be missing many of the same genes [cox3, atp6, atp8, nad6, and most tRNA genes]. One striking possibility is that these genes are present but cannot be found due to sequence divergence from the generalist ‘Protozoan mitochondrial DNA code’ that models like Mfannot, Mitofinder, and MITOS2 employ. Future studies that use long-read next-generation sequencing will hopefully help to clarify this discrepancy.

The identified *Rhodospirillales* bacterium potentially appears to be part of a new family, which clusters between the families *Thalassobaculaceae* and *Aestuariispiraceae* based on the phylogenetic tree assembled from multiple gene protein sequence alignment (Extended Data Fig. 7). Since this bacterium has not been isolated or cultured yet, we will not propose any nomenclature for it in this study. We simply provide a description of its annotated genome and propose its place amongst *Rhodospirillales*. The *Stentor hondawara* genome presented here is a high-quality (182.7 N’s per 100 kbp), very high coverage (100-500x), contig-level genome. Our initial assembly with SPAdes did not generate larger scaffolds, as we use exclusively short paired-read sequencing for this assembly. We attempted to generate scaffolds from the ∼3700 contigs using RagTag^54^ and the *S. pyriformis* genome as a reference, but this resulted in no quality scaffold generation, likely because the two *Stentor* species are very different at the DNA sequence level. This apparent issue also prevented a successful reference-guided genome assembly with MaSuRCA, though reference-guided assembly can work when assembling the genome of a *Stentor* species more closely related to either of the available reference genomes. Currently, *Stentor* genomes are assembled from the macronucleus, and ciliate macronuclear genomes are often fragmented down as small as single genes and subject to significant rearrangement^55–57^. Therefore, it is not surprising that no *Stentor* genome has achieved beyond scaffold-level contiguity. Indeed, even the *S. pyriformis* genome, which was assembled with PacBio long reads, could only achieve scaffold-level assembly (∼200 contigs) as it is an assembly of the macronuclear genome. It is currently unfeasible to extract enough micronuclear material from *Stentor* to generate a higher quality genome assembly.

Though we were only able to provide functional annotation for 21% of all 20,565 predicted protein-coding genes of the *Stentor hondawara* genome, these annotations include many important eukaryotic gene groups (Fig. 4a), and still revealed significant differences in the enrichment of genes between our unique marine *Stentor* species and the two previously annotated freshwater species (Fig. 4-5). Intriguingly, many enriched gene groups gave hits to plant circadian-related genes, including genes involved in seed dormancy, flowering timing, blue light response, and low fluence light response (Fig. 4a-b). These were enriched in *Stentor hondawara* over both freshwater *Stentor*, though in comparison to *S. coeruleus* alone, these were not enriched. Further inspection revealed that similar genes are enriched in *S. coeruleus* over *S. pyriformis* – explaining why these enrichments were missing from the “freshwater enriched genes” as this list included only genes enriched in both freshwater species. This enrichment, present in both *Stentor hondawara* and *S. coeruleus*, is interesting as such genes could likely be involved in the regulation of the *Stentor* circadian clock. The lack of these gene enrichments in *S. pyriformis* could be due to the presence of an obligate endosymbiotic *Chlorella variabilis* microalgae, which likely provides partial or full regulation of the circadian clock in this *Stentor* species^58^.

Among gene enrichments, marine enrichment for potassium and chlorine ion transport compared to freshwater stentor enrichment for sodium ion transport could be of interest, but requires further investigation. Many of the enriched marine genes (Fig. 5a-b ‘Metabolism’) are part of the biosynthesis and metabolism of various amino acids including alanine, glutamate/glutamine, aspartate, and glycine (Extended Data Table. 1). Interestingly, several amino acids, including primarily proline but also alanine and glutamate/glutamine, asparagine, and arginine, have been implicated as potential osmoprotectants in plants and other organisms^59–61^. It is therefore possible that the marine-specific enrichment for genes involved in the metabolic processes of alanine, asparagine, and glutamate/glutamine could be due to a role for these amino acids as osmoprotectants in *Stentor hondawara*. Consistent with this idea, while there are hits to alanine and glutamate/glutamine among the freshwater-enriched genes (Extended Data Table. 2), these are single hits compared to two hits for alanine and five for glutamate/glutamine in *Stentor hondawara*. Another interesting enrichment, which could warrant further investigation, is enrichment in the genes encoding static microtubule bundle and protein localization to the cell cortex in *Stentor hondawara* over freshwater stentors (Fig. 5b ‘Cell Architecture’). Taken together, these comparative genomics data suggest that the marine stentor, *Stentor hondawara*, enriches genes that help it manage water balance and survive environmental processes by regulating osmotic pressure and protecting cellular structures.

Aquaporins are important osmoregulators that allow exchange of water (orthodox aquaporins) and other small molecules like glycerol and ammonia (aquaglyceroporins). Our analysis of the aquaporins present in *Stentor hondawara*, along with *S. pyriformis* and *S. coeruleus*, revealed a large family of unique aquaporins unlike the aquaporins of more modern eukaryotes (plants, animals, and fungi). Interestingly, a subset of *Stentor* aquaporins, which was enriched (possibly through two duplication events) in *Stentor hondawara*, appears to be structurally – and thus potentially functionally – similar to aquaglyceroporins from other eukaryotes. Future validation of this possibility will require identification of the Ar/R selectivity filter, which generally consists of an aromatic and an arginine in close proximity within the aquaglyceroporin channel^62^. While the overall number of aquaporins present in each stentor does not vary significantly, this enrichment for a subset of potentially specialized aquaporins could belie a unique adaptation required for marine life. Future work could look at the knockdown of this subset of aquaporins in *Stentor hondawara* to test for sensitization to high salinity marine water. Similar enrichment and diversity of aquaporins and aquaglyceroporins and their link to seawater adaptation mechanisms have been identified in the fishes traveling across oceans and rivers, such as eels and salmon^63–65^. Furthermore, the other vertebrates, such as frogs, utilize aquaporins for aquatic-terrestrial adaptations, and use aquaglyceroporins to regulate glycerols as osmoprotectants for cold/freeze tolerance^66–68^. These cross-species findings, along with our study, suggest the ancestral and critical roles of aquaporins and aquaglyceroporins in the unique aquatic adaptations across the animal kingdom.

The *Stentor hondawara* mitochondrial genome is strikingly AT-rich with an average GC content of 13% (Fig. 3). Despite its small size of ∼48 kbp compared to the published *S. coeruleus* ∼300 kbp mitochondrial genome, the *Stentor hondawara* mitochondrial genome contains 42 genes, including 27 protein-coding genes and several rRNA and tRNA genes, similar to other complete mitochondrial genomes^69^. Furthermore, this was the single contig with ∼1000x coverage and such low GC content. We are therefore confident that this lone contig consists of the complete mitochondrial genome. Such compactness is not entirely unexpected, as some animal mitochondrial genomes can be as small as 16 kbp^69,70^. An unpublished genome assembly of a local *Stentor* isolated from the Virginia Tech campus, which is morphologically indistinguishable from *S. coeruleus*, has an identified mitochondrial genome of approximately 42 kbp in size, much closer in scale to the mitochondrial genome we include in this work.

One intriguing component of the hologenome is the presence of the genome assembly of a bacterial species in the order *Rhodospirillales* that is approximately 2.3 Mbp with ∼200-400x coverage and GC content of 43.37%. By all of these metrics, this assembly is high quality with coverage similar to the *Stentor hondawara* genome but with a significantly different GC content profile. This points to the possibility that the bacterial genome assembly is likely endosymbiotic rather than an environmental contaminant or food source. Several *Alphaproteobacteria*, including *Rhodospirillales*, are common endosymbionts within protists and other microbial marine eukaryotes. Indeed, some theories on the evolution of mitochondria suggest *Rhodospirillales* or *Rickettsiales*, a sister order in the class *Alphaproteobacteria*, as the potential ancestors of the mitochondrion^71,72^. Our potential endosymbiotic species appears to be a nutritional factory for the host cells, as it contains protein-coding genes that make it competent at nitrogen fixation, vitamin B12 production, production of other less complex B vitamins, amino acid biosynthesis, respiration, and energy production. Interestingly, several key genes for protection against foreign/phage DNA and cell wall synthesis are also present, while others, such as biofilm formation, quorum sensing, and flagellar assembly, are missing or fragmented. This finding, along with the intermediate size of the genome – 2.3 Mbp compared to ∼4-5 Mbp for intact *Rhodospirillales* genomes and ∼1 Mbp for more ancient endosymbionts – suggests that this organism is a recently acquired endosymbiont in the evolutionary history of *Stentor hondawara.* Strikingly, *Stentor hondawara* appears to have over two dozen isoforms of an ortholog to ABCD4, the transporter responsible for moving vitamin B12 from lysosomes into the cytosol in eukaryotes, compared to approximately 4 in humans. This enrichment suggests that *Stentor hondawara* has specialized in receiving B12, perhaps from its endosymbiotic bacteria. A final piece of evidence that is missing from our analysis of whether this bacterial species is an endosymbiont or not is demonstrating that we can extract it from the stentor cells and identify it as an *Alphaproteobacteria* from this source. Though we did not have enough live samples to attempt this, we would point out that most *Stentor hondawara* cells we imaged presented with multiple patches of rust-colored to reddish-brown coloration, which could be clusters of the *Rhodospirillales* bacteria within the stentor.

In stark contrast to the strong signal of our putative endosymbiotic bacteria, several other *Pseudomonadota* and *Chlorophyta* contigs with low coverage (5-50x) were also present in the assembly and were removed from the hologenome as likely contaminants from the culture water or partly-digested food from the stentor cells. One strong candidate as a potential food source is *Bathicoccus prasinos*, which was initially identified in an analysis of the raw genome assembly as a 0.01% contaminant (1% of all contaminant DNA). *Bathicoccus prasinos* is a marine picoplankton that is ubiquitous in marine environments, making it a likely food for *Stentor hondawara*. As all long-term culturing efforts have failed to maintain stable cultures beyond a month after leaving the source of fresh Cape Cod ocean water, likely due to a failure to identify a stable food source, we will pursue this and other picoplankton as a potential food for culturing. Several strains of *Bathicoccus prasinos* and other potential food microbes can be obtained from online repositories like the UTEX Culture Collection (https://utex.org/) and can be cultured in a lab setting with minimal difficulty. These picoplankton and other microbes will be explored as potential food sources for future *Stentor hondawara* culturing efforts.

As future efforts reveal more freshwater and marine *Stentor* species, it will be interesting to perform a more exhaustive comparison of enriched genes between the two vastly different types of environments. Furthermore, a direct comparison between a marine species and a close freshwater relative could be particularly informative. In the case of *Stentor hondawara*, its closest relative is currently *S. katashimai*, a freshwater species found in 1973 in Japan^12,13^, which has not been seen since it was last isolated and had its 18S sequence published in Germany in 2010^73^. Once *S. katashimai* is reisolated or a closer relative discovered in the near future, its whole-genome will be of special importance to compare with *Stentor hondawara* on the in-depth identification of an essential set of genes for marine adaptation.

Overall, our discovery of the first verified marine stentor species, *Stentor hondawara*, and the comparative genomics we performed against two well-established freshwater *Stentor* species, opens a new avenue for the exploration of *Stentor* species, including both marine and freshwater, and provides insights into the evolutionary and minimal mechanisms of environmental adaptation employed at the scale of single-cells.

## Methods

### Collection

Marine water samples were collected daily on clear sunny days, usually within two hours of high tide in the afternoon, from early July through mid-August in 2023, 2024, and 2025. Some samples were also collected within 2 hours of low tide, specifically in 2025. Collection was performed with a combination of a 50-micron mesh plankton net (Carolina Biological) and a plastic beaker attached to a retractable handle. At each collection, the plankton net was towed up and down the jetty behind the NOAA parking lot on the MBL campus (41°31’29.8" N 70°40’26.9" W), keeping the net about one to five feet below the surface so as to concentrate near-surface plankton and other microbes. Following one or two passes up and down the jetty, the concentrated microbes were collected in a 10-gallon bucket half-filled with ocean water. Next, the plastic beaker was used to collect smaller volume surface water samples, focusing specifically on collecting small clusters of the brown kelp, *Sargassum,* floating in the vicinity of the jetty (see also ‘Etymology’). All samples were added to the same 10-gallon bucket and taken back to the lab for isolation. Overall, we collected about 800 live *Stentor hondawara* individuals in this study. Of these, 300 non-clonal cells were used for the DNA extraction (see also ‘DNA extraction’).

### Isolation

Marine samples were processed by taking a small aliquot into a 100 mm deep dissection petri dish (approximately 100 ml of water at a time) and placing the sample on a stereomicroscope. While looking through the microscope, individual *Stentor hondawara* cells were isolated with a 200 ul pipette and placed into a sample petri dish (60 mm) with filtered Pacific Ocean Water (fPOW; at 35 ppt salinity; Imagitarium). Any organisms larger than the stentor cells were removed from the isolated stentor cultures. Isolation was primarily performed in the evenings as *Stentor hondawara* cells were more motile at this time.

### Culturing

Isolated marine stentors were kept in either 6-well plates, 60 mm petri dishes, or 100 ml wide-mouth glass jars filled with either Imagitarium brand fPOW mixed with partially filtered source ocean water (from initial collection; filtered for larger organisms), or solely in partially filtered source ocean water. Some cultures were supplemented with commercially available phytoplankton (Seachem phytoplankton and AlgaeBarn phytoplankton mixes); these required water changes every 3 to 4 days. Water changes entailed replacing half of the water with fresh purified fPOW or with fresh ocean water from the source. Fresh water was partially filtered by manually removing organisms larger than *Stentor hondawara* cells before being introduced to cultures. The food organism(s) in this water are as yet unknown. All cultures were kept between 18-20 °C and were kept under natural light near a windowsill (approximately 13.5-14.5 hours of natural sunlight daily in the summer months). To date, we have been unable to maintain healthy cultures beyond a few weeks away from the MBL.

### Microalgae culturing

Mixed phytoplankton cultures from commercial sources were grown in makeshift bubbler columns full of ocean water with a large bubbler stone connected to an air pump generating enough bubbles to consistently mix the water column and keep the algae in suspension. Cultures were kept under very bright white light (200 lumens) for 18 hours a day. Every two weeks, water was supplemented with liquid fertilizer (MiracleGro) to add back nutrients. Salinity was regularly monitored with a refractometer, and distilled water was added back to stave off evaporation and keep salinity at around 35 ppt, corresponding to the average salinity of ocean water.

### DNA extraction

Three hundred non-clonal *Stentor hondawara* cells were isolated in fPOW and starved for 72 hours. DNA extraction followed a standard phenol-chloroform extraction protocol as follows. 300 starved cells were collected into a 1.5 mL centrifuge tube, centrifuged at 5,000 g for 2 minutes, and all liquid was removed and replaced with 200 µl of PBS. The 200 µl cell suspension was then heated to 65 °C for 10 minutes and then vortexed twice for 15 seconds to homogenize the partially lysed cells. Next, 200 µl of phenol/chloroform/isoamyl alcohol (25:24:1; pH 8.2) solution was added, and the sample was vortexed for 1 minute, after which the sample was centrifuged at maximum speed (minimum of 12,000 g) for 5 minutes. Approximately 180 µl of the top aqueous DNA solution was removed and placed into a new tube, taking care not to disturb the bottom phenol/chloroform/isoamyl alcohol layer. 200 µl of elution buffer (Qiagen recipe) were added to the tube with phenol/chloroform/isoamyl alcohol, and the mixture was vortexed for 1 minute and centrifuged at top speed for 5 minutes again. Once more, approximately 180 µl of the top aqueous layer was moved to the same new tube as before. Next, 360 µl of chloroform/isoamyl alcohol was added to the ∼360 µl of DNA solution, the sample was vortexed for 1 minute and then spun in a centrifuge at maximum speed for 5 minutes. Approximately 360 uL of the top aqueous solution was removed and placed into a new tube, taking care not to disturb the bottom chloroform/isoamyl alcohol layer. Next, 40 µl of 7.5 M NH4OAc and 1 µl of glycogen (20 µg) were added to the tube with the DNA sample. After mixing via vortexing for 10 seconds, 1,000 µl of 100% ethanol was added to the tube, and the sample was placed at -20 °C for 8 hours. Next, the sample was centrifuged at maximum speed for 20 minutes at 4 °C, and the supernatant was carefully decanted without disturbing the DNA pellet. The DNA pellet was next resuspended in 300 µl of 80% ethanol and vortexed 3 times for 15 seconds each. The sample was then centrifuged at maximum speed for 15 minutes at 4 °C, and the supernatant was decanted without disturbing the pellet. The 80% ethanol wash and decanting were repeated a second time, after which a final centrifugation at top speed for 30 seconds was performed. Residual ethanol was removed carefully, and the pellet was air-dried for 30 minutes at room temperature. The DNA was resuspended in 40 µl of DNAse-free water. The final sample was checked on a Nanodrop for DNA quality and concentration, and stored at -80 °C until use for genome sequencing. A portion of this initial sample remains in a -80 °C freezer at MIT.

### Whole-genome sequencing and raw assembly of *Stentor hondawara* genomic DNA

50 ng of the extracted *Stentor hondawara* genomic DNA (see ‘DNA extraction’) was prepared for short read sequencing using Illumina Nextera Flex (Illumina) chemistry using modified oligonucleotides for amplification, replacing the Illumina P5/P7 anchor sequences with Singular S1/S2. Libraries were amplified for 15 cycles, quantified by qPCR and on an Agilent Fragment Analyzer, and sequenced on a Singular G4 sequencer (Singular Genomics) using 1 lane of an F3 flowcell run at 150 nt paired end. *De novo* assembly and rRNA gene prediction were performed using the nf-core/mag pipeline^17^. This pipeline was used to run *de novo* assemblies of quality-checked (FastQC) and trimmed (Trimmomatic) paired reads with SPAdes (v4.1.0)^18^ and MEGAHIT (v1.2.9)^74,75^, along with additional annotation and quality-control modules, including protein predictions from MEGAHIT which we did not use, but we instead used GeMoMa^14,15,22^ for protein predictions using the gene annotations of *S. coeruleus*^6^ and *S. pyriformis*^7^. We also attempted a referenced genome assembly using MaSuRCA^19^. To select which genome to use as a reference, we first used Bowtie2 to map our raw reads to both *S. pyriformis* and *S. coeruleus* published genomes. Bowtie2 revealed very low alignment to either genome, but with 10-fold higher overall mapping to the *S. coeruleus* genome (4% alignment vs 0.6% alignment). We therefore used the *S. coeruleus* genome as a reference for MaSuRCA assembly, using 150 bp overlaps and 350 bp average read lengths, which we calculated from the raw read files using SAMtools. Despite running for 96 hours with 64 CPUs and 650 GB of memory allotted to it on a dedicated computing cluster, MaSuRCA did not yield a viable assembly and gave an initial estimate of 169 Mbp, significantly over-estimating the actual genome size. QUAST (v.5.3)^76,77^ was used to analyze the quality of the two *de novo* genome assemblies generated by nf-core/mag, and the highest quality assembly, the SPAdes assembly in our case, was retained for further processing. This assembly was 98.3 Mbp with an N50 of 55.1 kbp and 149.78 N’s per 100 kbp.

### Genome cleanup and separation

The initial genome assembly was used to predict proteins with GeMoMa^14,15,21^ using the gene annotations of *S. coeruleus*^6^ and *S. pyriformis*^7^, both available from the *Stentor* database (stentor.ciliate.org), as references for protein-coding gene prediction. This initial annotation was then run through a BLASTp DIAMOND^24^ search against the nr_2023-09-03 reference database, using the 2024-06-05 NCBI taxonomy database with “very-sensitive” sensitivity and BLOSUM45 protein alignment matrix, on Galaxy (usegalaxy.org). We also generated a megablast (BLASTn) against the raw genome assembly using the nt_2023-09-01 database – though this yielded low quality hits and was not used for analysis. Finally, we also ran a BUSCO ortholog analysis on the genome assembly and generated a coverage map of the raw genome assembly using Minimap2^78^ and the raw reads. All of these analyses, except the megablast results, were pooled into BlobToolKit^79^ in order to visualize the genome contigs along with their GC content, coverage, and identified taxonomy IDs (NCBI taxonomy). Using Blobtools2^79^, we separated the *Stentor* genome from contaminants by GC content (∼30% documented GC content for *S. pyriformis* and *S. coeruleus*). Most contigs identified as *Ciliate* demonstrated coverage between 100x and 500x, we therefore chose to filter out any contigs below 100x coverage and with fewer than 500 base pairs (to filter out fragments smaller than single genes).

A significant cluster of *Rhodospirillales* was identified by BlobTools2. Based on coverage and GC content, we determined this to be a possible endosymbiont and filtered it out from the final *Stentor* genome assembly (for this assembly, we included contigs of length above 200 bp) along with several “No-hit” contigs that clustered tightly with the *Rhodospirillales* cluster. Any “No-hit” contigs that contained greater than 50% *Ciliophora* hits were returned to the *Stentor hondawara* assembly; all other contigs were kept as part of the *Rhodospirillales* genome. This set of contigs was annotated using Prokka^38^ and BLASTx databases on Galaxy. BUSCO analysis of this genome was run using the rhodospirillales_odb12 dataset.

We also separated out the putative mitochondrial genome of *Stentor hondawara*, found as a single contig (NODE_319) based on its low GC content and high coverage (several fold higher than the *Stentor hondawara* genome contigs). We verified this contig as the mitochondrial genome by identifying and annotating mitochondrial genes using Mfannot^20^ and tRNA gene using barrnap^21^ (Extended Data Table. 3). We also tried MitoFinder^80^ running MEGAHIT^75^, metaSPAdes^81^, and IDBA-UB^82^ assembly methods with the few *Ciliate* annotated mitochondrial genomes available in the NCBI database (no annotated *Stentor* mitochondrial genome has been posted) but none yielded any mitochondrial gene hits. The mitochondrial genome, as annotated by Mfannot, appeared to be missing several key genes, including cox3. We therefore also tried *de novo* annotation with MITOS2^83^, which yielded more complete annotations, but still lacked several key genes, including cox3. As a control, we also annotated the published *S. coeruleus* mitochondrial genome using MITOS2 and found a nearly identical group of genes missing. The mitochondrial genome, in its current state, has been uploaded to GenBank (submitted and awaiting Accession ID).

### 18S SSU alignment and *Stentor* phylogenetic tree assembly

A local BLAST genomic database was assembled using the decontaminated genome assembly of *Stentor hondawara*. The complete 18S SSU sequence of *S. pyriformis* (GenBank Accession ID: LC533385.1) was used to run a BLASTn search against the *Stentor hondawara* genome database, resulting in a couple of hits with high sequence conservation. The strongest hit, a contig 4kbp in length, gave the reverse complement of the *S. pyriformis* full sequence. This contig, after being reversed, was aligned with the *S. pyriformis* sequence using MAFFT^34^ (200PAM nucleotide alignment matrix), and all non-overlapping regions were trimmed off. The resultant sequence was 2,044 base pairs long and was submitted to GenBank (Accession ID: PX841742). Reference sequences were assembled from GenBank for all available *Stentor* species, along with *Maristentor dinoferus* (Accession ID: AY630405.1), as *Stentor hondawara* could have been a novel species of genus *Maristentor*, *Blepharisma americanum* (Accession ID: KP970223.1) as an outgroup, and *Condylostentor auriculatus* (Accession ID: KP970235.1) as the root node. All sequences were aligned using MAFFT (200PAM). Resultant alignments were trimmed in UGENE, keeping only approximately 560 base pairs of full overlap common to all sequences. Trimmed sequences were analyzed using IQTree v3^84^ to infer a maximum-likelihood (ML) tree that was rooted using *Condylostentor auriculatus, Maristentor dinoferus,* and *Blepharisma americanum* as the outgroups. The ‘automatic’ substitution model method was used to identify the optimal nucleotide substitution model, HKY+I+G4. Branch reliability was assessed using bootstrapping with 10,000 replicates as well as 1000 replicates of single-branch Shimodaira-Hasegawa approximate likelihood ratio test (SH-aLRT)^85^ and single-branch Bayesian-like transformation of aLRT (aBayes)^86^. The resultant tree was visualized using FigTree (v1.4.5) and saved as an SVG file for figure generation.

### Multiple protein alignment and *Rhodospirillales* phylogenetic tree assembly

Three core vitamin B12 biosynthesis pathway proteins [CobN, CobT, CobS] and another core protein [SecD] were used to generate alignments for this phylogenetic tree. BLASTp was used to find the respective four proteins from members of each major *Rhodospirillales* family/cluster, along with a representative of *Rickettsiales* as an outgroup. These were concatenated into a single ‘sequence’ for each query and then aligned using MAFFT (BLOSUM62 protein alignment matrix), and all non-overlapping regions were trimmed off using UGENE. Trimmed sequences were analyzed using IQTree v3 to infer a maximum-likelihood (ML) tree based on amino acid sequence identity. The ‘automatic’ substitution model method was used to identify the optimal substitution model, LG+F+I+G4. Branch reliability was assessed using bootstrapping with 10,000 replicates as well as 1000 replicates of single-branch Shimodaira-Hasegawa approximate likelihood ratio test (SH-aLRT) and single-branch Bayesian-like transformation of aLRT (aBayes). The resultant tree was visualized using FigTree (v1.4.5) and saved as an SVG file for figure generation.

### Comparative genomics with gene annotations

The decontaminated *Stentor hondawara* genome was then soft-masked with RepeatMasker^87^ using a model of low-complexity repeats trained on the *Stentor hondawara* genome using RepeatModeler^88^. The final soft-masked genome was then run through GeMoMa once again using the structural annotations of *S. coeruleus*^6^ and *S. pyriformis*^7^, both available from the *Stentor* database (stentor.ciliate.org), as references for gene prediction. The GeMoMa annotation was then analyzed for completeness by comparing its BUSCO metrics to the metrics of the raw *Stentor hondawara* genome, and the metrics of the annotations of *S. coeruleus* and *S. pyriformis.* Next, OrthoFinder^26^ was run on all three protein annotations (our GeMoMa prediction and the two published assemblies) in order to generate orthogroups common amongst all three and unique to each. Following orthogroup identification, eggNOG (v5.0.2; https://eggnog5.embl.de/)^23,24^ was used to generate a partial functional GO annotation of genes for all three species individually, using their individual protein sequence read files, with ultra-sensitive Diamond settings and with the BLOSUM45 protein alignment matrix. BLOSUM45 was selected as it is more likely to find more divergent hits, which could be especially useful given that the *Stentor* genus is very divergent from most other Eukaryotes represented in the eggNOG database. Finally, topGO^89^ (R package) was used to analyze orthogroups specific to *Stentor hondawara* or to the freshwater species, for enrichment and to provide functional annotations where available. Weighted Fisher was used to rank enrichment, with a minimum node size of 4 (the minimum number of genes in the GO group). All enriched genes with a score below 0.05 generated for BP, MF, and CC categories were combined into a table for either *Stentor hondawara* only, or both freshwater species (*S. coeruleus* and *S. pyriformis*) together only.

### Protein structure prediction

Proteins of interest were identified by BLASTp against known human orthologs. The top identified *Stentor hondawara* orthologs were isolated in FASTA format and fed into Alphafold 3 through the Alphafold server (https://alphafoldserver.com/). The top protein structural prediction model for each was downloaded as a pdb file and visualized using PyMol (v3.1.6.1). Stentor orthologs were aligned to human orthologs using the ‘super’ command; resultant RMSD statistics were recorded and reported above.

### Aquaporin phylogenetic tree assembly and ProtSpace clustering

All identified *Stentor hondawara* aquaporin protein sequences were pooled together with all aquaporins identified in *S. pyriformis* and *S. coeruleus* genomes, all human aquaporins, yeast aquaporins (*Saccharomyces cerevisiae*), algal aquaporin (*Bathicoccus prasinos*), major nematode aquaglyceroporins (*Caenorhabditis elegans*), and plant aquaporins (*Arabidopsis thaliana*). The joint FASTA file was first aligned using MAFFT (BLOSUM30) and used to generate a protein sequence-based ML tree with IQTree v3 using the Q.PFAM+I+G4 model (selected by IQTree best model selection). The resultant tree was visualized with FigTree v1.4.5 and exported as an SVG file for final figure assembly in Adobe Illustrator. The unaligned FASTA sequences were also fed into the ProtSpace Google Colab notebook designed for protein embedding (https://github.com/tsenoner/ProtSpace)^35^ to generate embedding and feature files. These were then fed into the visualization Google Colab notebook. 2D-clustering was performed by t-distributed Stochastic Neighbor Embedding (t-SNE) dimensional reduction using a perplexity value of 32. The resultant 2D plot was saved as a PNG file for figure generation.

### ESPript assembly and Aquaglyceroporin functional domain conserved sequence comparison

Aligned Aquaglyceroporin sequences were separated into a discrete CLUSTAL (.aln) file and plotted against human AQP3 (prototypical Aquaglyceroporin) using ESPripr v3.2 (online tool; https://espript.ibcp.fr/ESPript/ESPript/)^90^ to plot residue alignment along with the predicted 2nd order structural elements of AQP3. Shown sequences highlight known conserved residues and functional domains that bear Aquaglyceroporin-specific modifications in many Aquaglyceroporins, including the FAT/FST motif, the two NPA motifs, and the N-terminal regulatory region where P2, P3, P4, and P5 reside^36,37^. Residue colors were set based on relative charge.

### Live cell imaging

Live cells were placed in the concave depression of a depression slide with fresh fPOW. Live imaging data were acquired on a Zeiss AxioZoom V.16 equipped with a PlanNeoFluar Z 2.3x/0.57 objective and a Thorlabs CS505CU 12-bit color CMOS camera (Thorlabs) or an Amscope stereomicroscope (SM2-TZ; AmScope) with a Thorlabs monochrome CMOS (CS135MUN; Thorlabs). Images and videos were acquired using the ThorCam software (Thorlabs) at approximately 20 frames per second. On either microscope, imaging was carried out using either direct transmitted light from below the stage or darkfield (oblique light) through a ring diffuser also below the stage. The scale for size measurements of live cells was established using a micrometer slide with 10-micron demarcations (AmScope), which was imaged at the same magnification as was used for the live imaging. The scientific sketch (Fig. 1d) was drawn using a Wacom Cintiq 16 pentablet (Wacom) with a Wacom Pro Pen 3.

### Fixation and fixed imaging

Cells were collected from sample cultures and washed in fresh ocean water (fPOW). 10-20 cells were then moved into ∼100 µl of fPOW atop a #1.5 22×22 mm coverslip, which was covered with a 1 mm thick silicone pad with a 10mm circular hole cut out of it. After cells were moved into the sample area of the silicone-padded coverslip, 100 µl of relaxation buffer [10 mM EGTA, 70 mM Tris, 3 mM MgSO4, 7.5 mM NH4Cl, 10 mM phosphate buffer (pH 7.1), pH to 7.2] was added and mixed with the fPOW. Cells were incubated for 10-15 minutes in the relaxation buffer, until cells were visibly extended. Following relaxation, most of the liquid was removed by pipette and 200 µl of 1% glutaraldehyde in PBS was added to the cells. After 60 minutes in glutaraldehyde, cells were washed twice, for 5 minutes each, with 200 µl of PBS. Next, PBS was removed and replaced with 200 µl of PBST (Tween-20) along with 1 µl of DAPI solution (20 ng/µl). After 5 minutes in PBST, most of the liquid was removed and replaced with ∼50 µl of Prolong Glass (Thermo Fisher) mounting media. Next, a slide was then inverted onto the coverslip, taking care not to trap any bubbles within the sample area. Excess liquid was wicked away, and the slide was left inverted overnight in a 4 °C refrigerator. After overnight drying, the slide was briefly turned right side up, sealed with nail polish, then re-inverted and stored with the coverslip down at 4 °C indefinitely or until imaging.

Imaging was carried out on two separate systems. For most basic imaging, an Olympus IX81 with a 40x 0.75 N.A. UPLAN FL N objective (Olympus) and a Thorcam CS126MU (Thorlabs) were used. For higher contrast imaging, some fixed samples were also imaged on a custom-built OI-DIC^16^ system running on an Olympus IX81 with a 40X 0.75 N.A. objective and a Photometrics camera (Teledyne) in the Shribak lab at the MBL.

## Data availability

We made the genomic sequences of *Stentor hondawara* (submitted and awaiting Accession ID) and its putative endosymbiotic bacteria (submitted and awaiting Accession ID) publicly available through the NCBI genome database. The *Stentor hondawara* mitochondrial genome and its annotation (by Mfannot) have been uploaded to GenBank. The other stentor species’ genome assemblies used in this study are also available on the DRYAD database through DOI: https://doi.org/10.5061/dryad.bzkh189kk (*S. coeruleus*) and through DOI: https://doi.org/10.5061/dryad.g79cnp612 (*S. pyriformis*).

## Acknowledgements

This study was supported by the Grass Fellowship (to T.H. and D.B.C.) funded by the Grass Foundation, the Whitman Fellowship (to T.H. and D.B.C.) funded by the MBL, and the Kavli-Grass-MBL Fellowship (to T.H.) funded by the Kavli Foundation, Alkermes Pathways Research Award (to T.H.), Osamu Hayaishi Memorial Scholarship for Study Abroad by the Japanese Biochemical Society (to T.H.), Uehara Memorial Foundation Overseas Fellowship (to T.H.), Japan Society for the Promotion of Science Overseas Research Fellowship (to T.H.), and Picower Fellowship (to T.H.) by the MIT Picower Institute and the Freedom Together Foundation.

We thank the MBL for laboratory space, technical and administrative support, especially Nipam Patel, Anne Sylvester, Briana Bertochi, Lisa Abbo, Marko Horb, and Michael Shribak. We also thank the administrative support, especially from Catherine Carr and Matthew McFarlane of the Grass Foundation, and from Amy Bernard and Angie Michaiel of the Kavli Foundation. We thank Cody Ayers for providing aerial drone footage of the collection area at the MBL. We appreciate guidance and advice on genomics and bioinformatics from Anton Suvorov at Virginia Tech, Mandië Holford at Harvard, and Huiming Ding, Charlie Whittaker, and Stuart Levine at MIT BioMicro Center and Barbara K. Ostrom (1978) Bioinformatics and Computing Core Facility of the Swanson Biotechnology Center. We appreciate Matthew Wilson at MIT for his generous support and guidance, and the Grass and Whitman Fellows for helpful discussion.

## Author Information

### Contributions

T.H. and D.B.C.: Conceptualization, Formal Analysis, Funding Acquisition, Investigation, Methodology, Supervision, Writing—original draft, Writing—review and editing.

### Corresponding authors

Takato Honda and Daniel B. Cortes.

## Extended Data

**Extended Data Figure 1.**
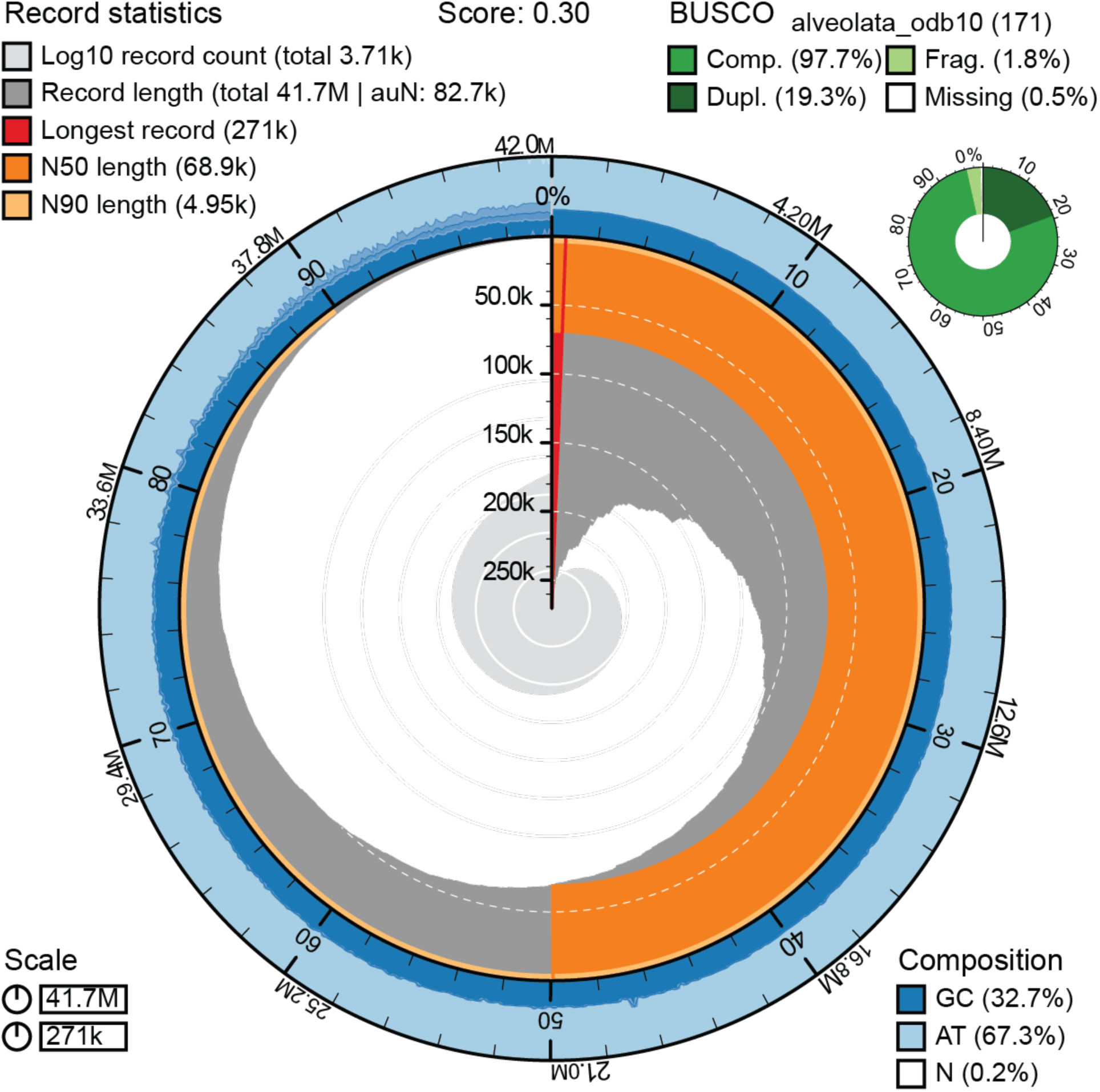
Metrics for the *Stentor hondawara* genome assembly. Snail plot showing the quality metrics for the genome assembly, including N50, GC content, total assembly size, Ns (as a percentage of total genome), and the initial BUSCO score for proteins predicted by GeMoMa from the raw genome assembly. The plot was made with Blobtools2.

**Extended Data Figure 2.**
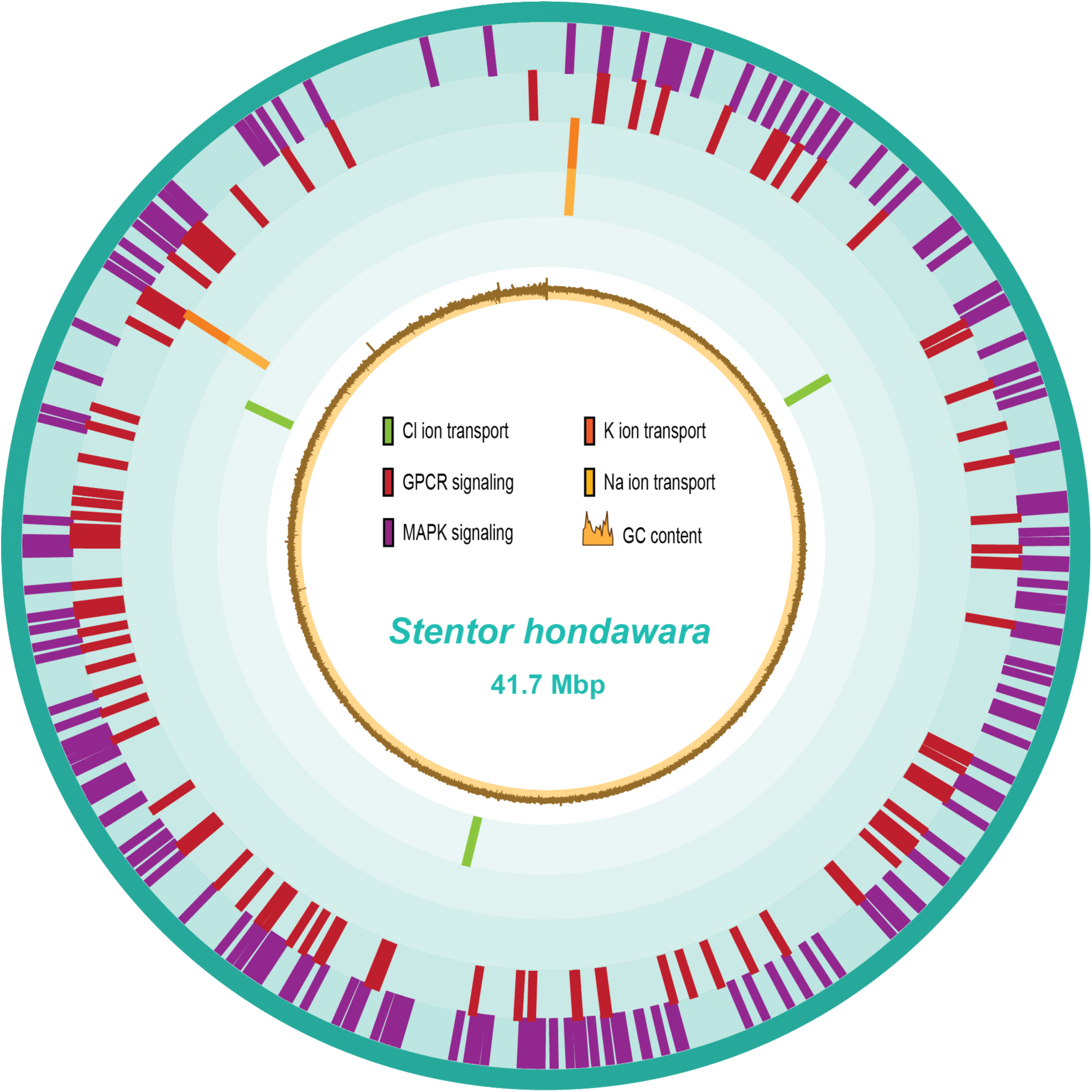
Graphical representation of *Stentor hondawara* genome. Graphical display of the whole *Stentor hondawara* genome, highlighting several functional groups of orthogenes that were uniquely enriched in *Stentor hondawara* over *S. pyriformis* or *S. coeruleus* (figure legend).

**Extended Data Figure 3.**
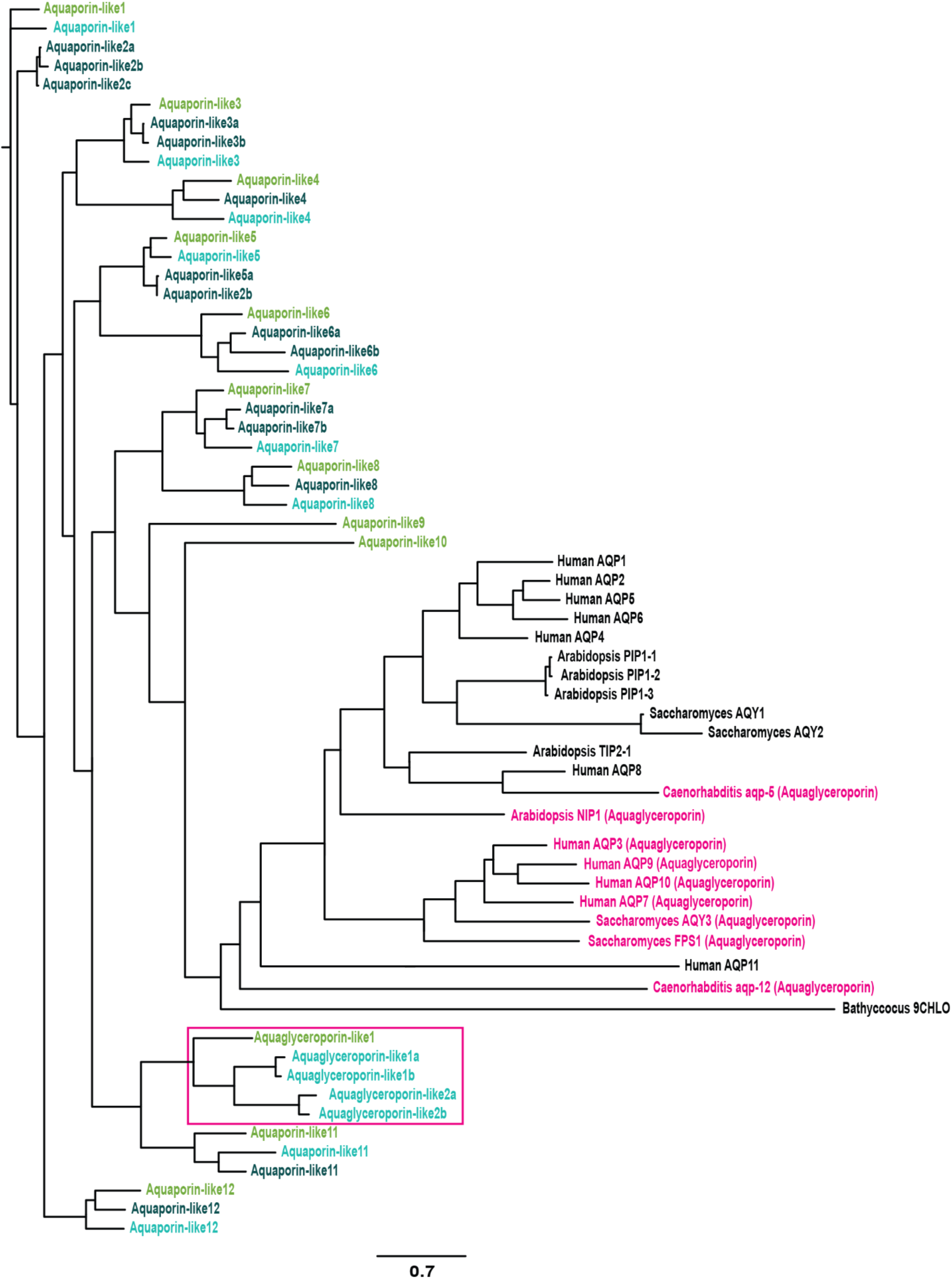
Phylogenetic tree of *Stentor* aquaporins compared to other Eukaryotic aquaporin families. Phylogenetic tree assembled using maximum likelihood (ML) estimates from peptide sequence identity, with branch lengths showing amino acid substitutions per site. *Stentor* families are color-coded for orthologs in *S. pyriformis*, *S. coeruleus,* and *Stentor hondawara.* Known aquaglyceroporins have pink text. Presumptive *Stentor* aquaglyceroporins are highlighted in a pink box. The scale bar shows substitutions per amino acid.

**Extended Data Figure 4.**
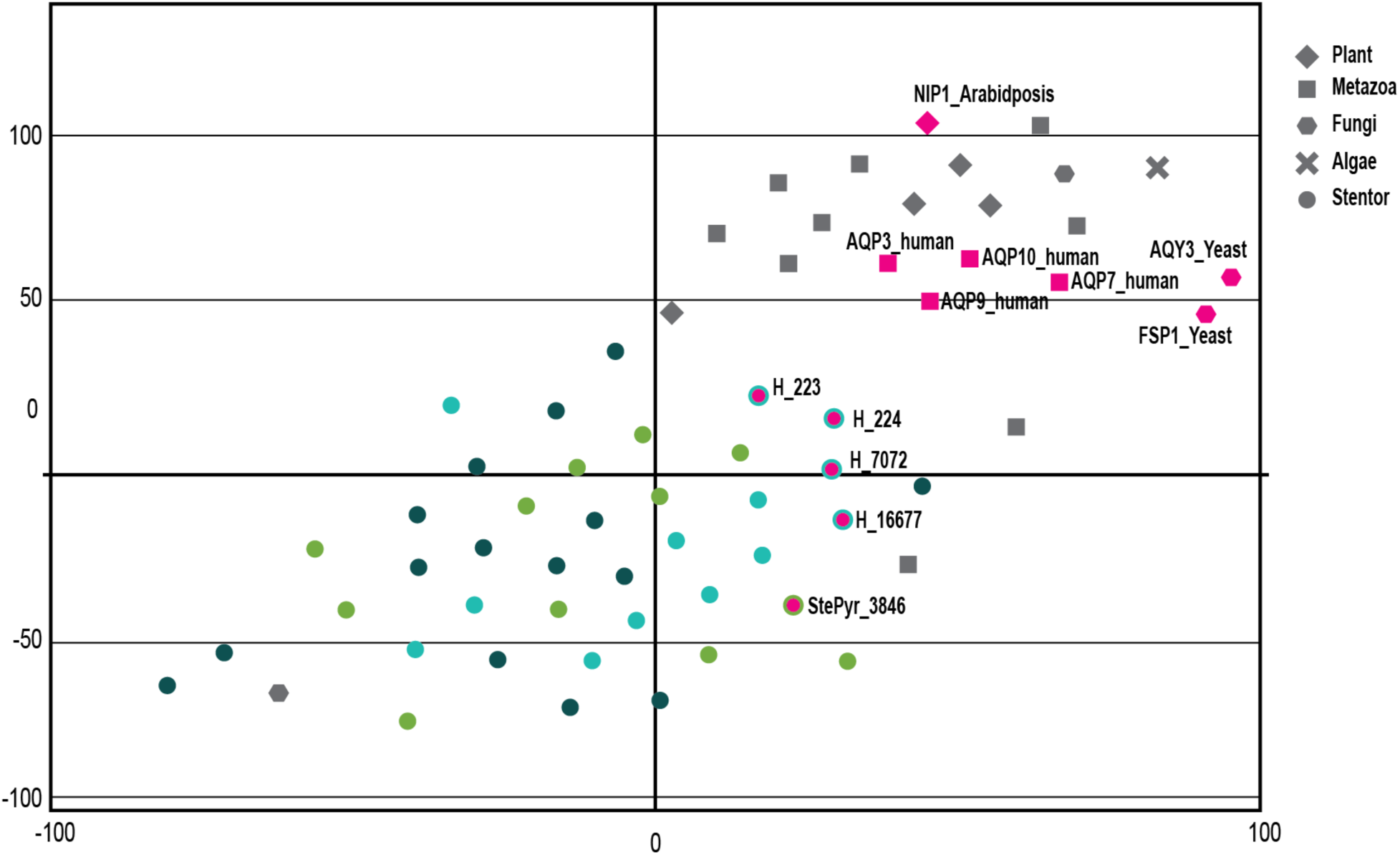
t-SNE map of aquaporin orthologs across Eukarya. 2D t-SNE cluster map showing the local and global structural similarity predicted based on protein sequence for all identified *Stentor hondawara* aquaporins along with human aquaporins (blue squares), human (pink squares), algal (teal diamond), and fungal (red X) aquaporin-7 orthologs. *Stentor hondawara* and *S. pyriformis* aquaporins closest in protein sequence to aquaporin-7 are highlighted as pink circles with border color specific to the species; all other “generic” *Stentor hondawara* aquaporins are shown as circles color-coded by species.

**Extended Data Figure 5.**
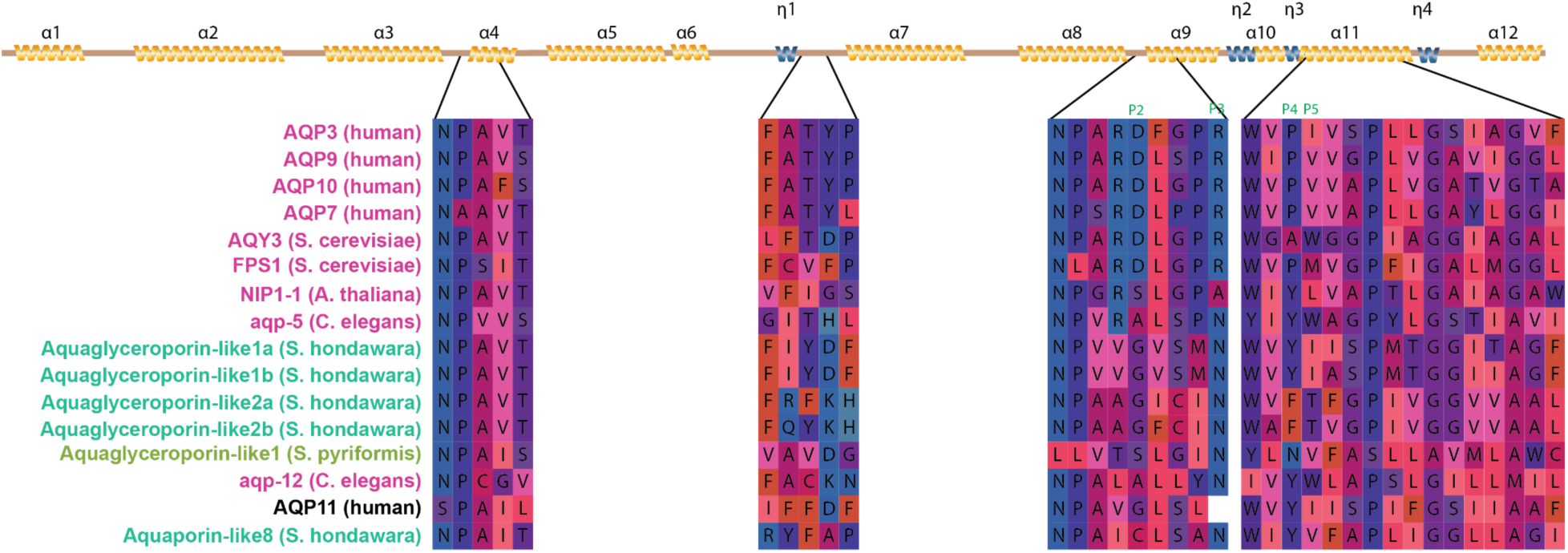
Comparison of conserved Aquaglyceroporin-specific domains between *Stentor hondawara* and other Eukaryotes. Conserved functional regions of Aquaglyceroporins aligned to human AQP3 (top sequence) secondary structure as assembled by ESPript 3.2. Colors are based on the relative charge of the residue. P2-P5 correspond to key residues identified for the classification of aquaglyceroporins. The bottom two sequences are generic aquaporins, for human (AQP11), and for *Stentor hondawara* (Aquaporin-like8).

**Extended Data Figure 6.**
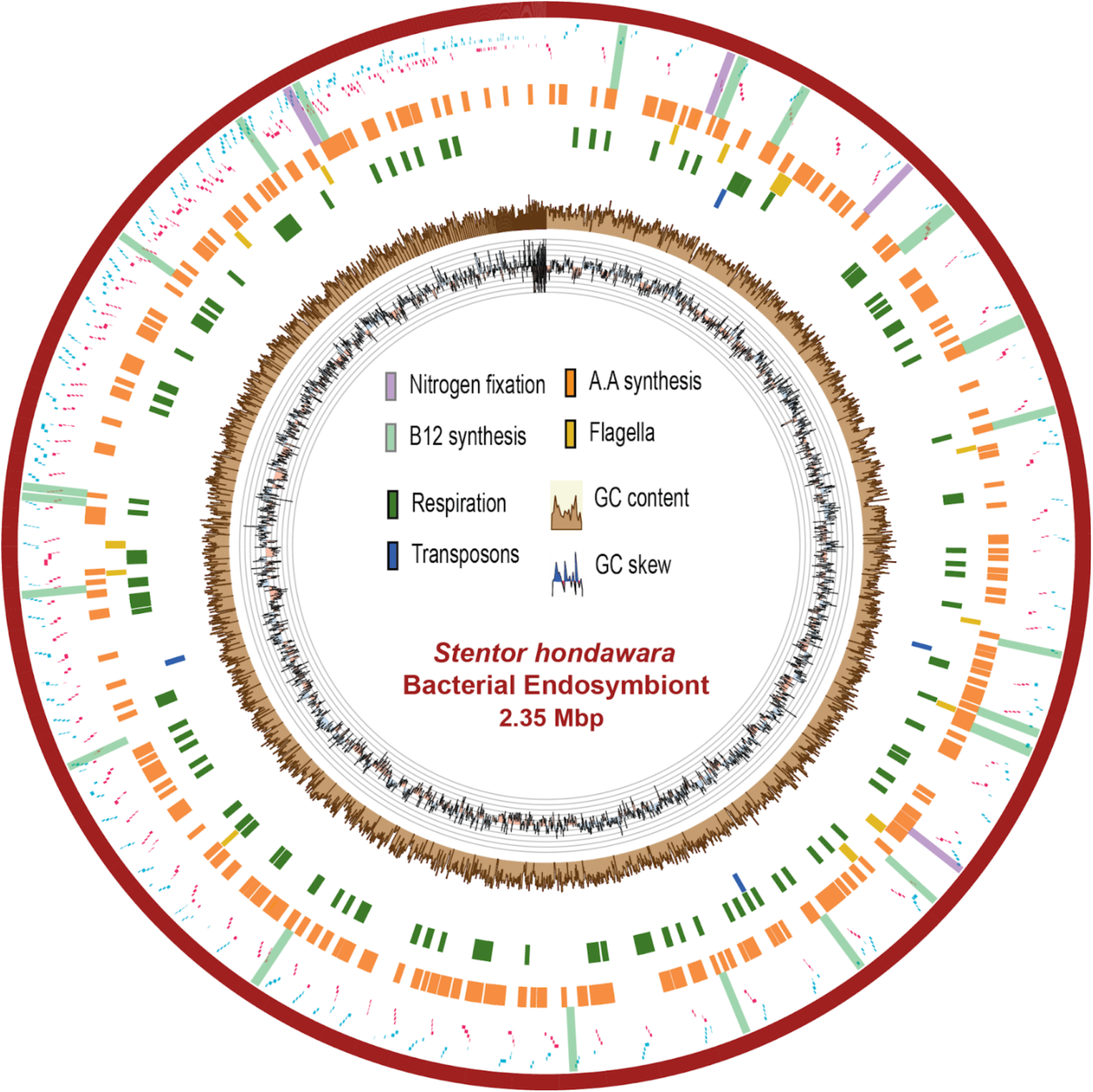
Graphical representation of the prospective endosymbiotic bacteria of *Stentor hondawara*. Graphical display of the fully assembled genome of the presumptive endosymbiotic bacteria found as part of the raw *Stentor hondawara* genome assembly. Small blue and red dots correspond to gene transcriptional start sites and whether they are on the positive (blue) or negative (red) DNA strand. All other highlights showcase functional gene groups of specific interest (figure legend). Two internal circular plots demonstrate overall GC content (outer brown plot) and GC skew (inner black plot).

**Extended Data Figure 7.**
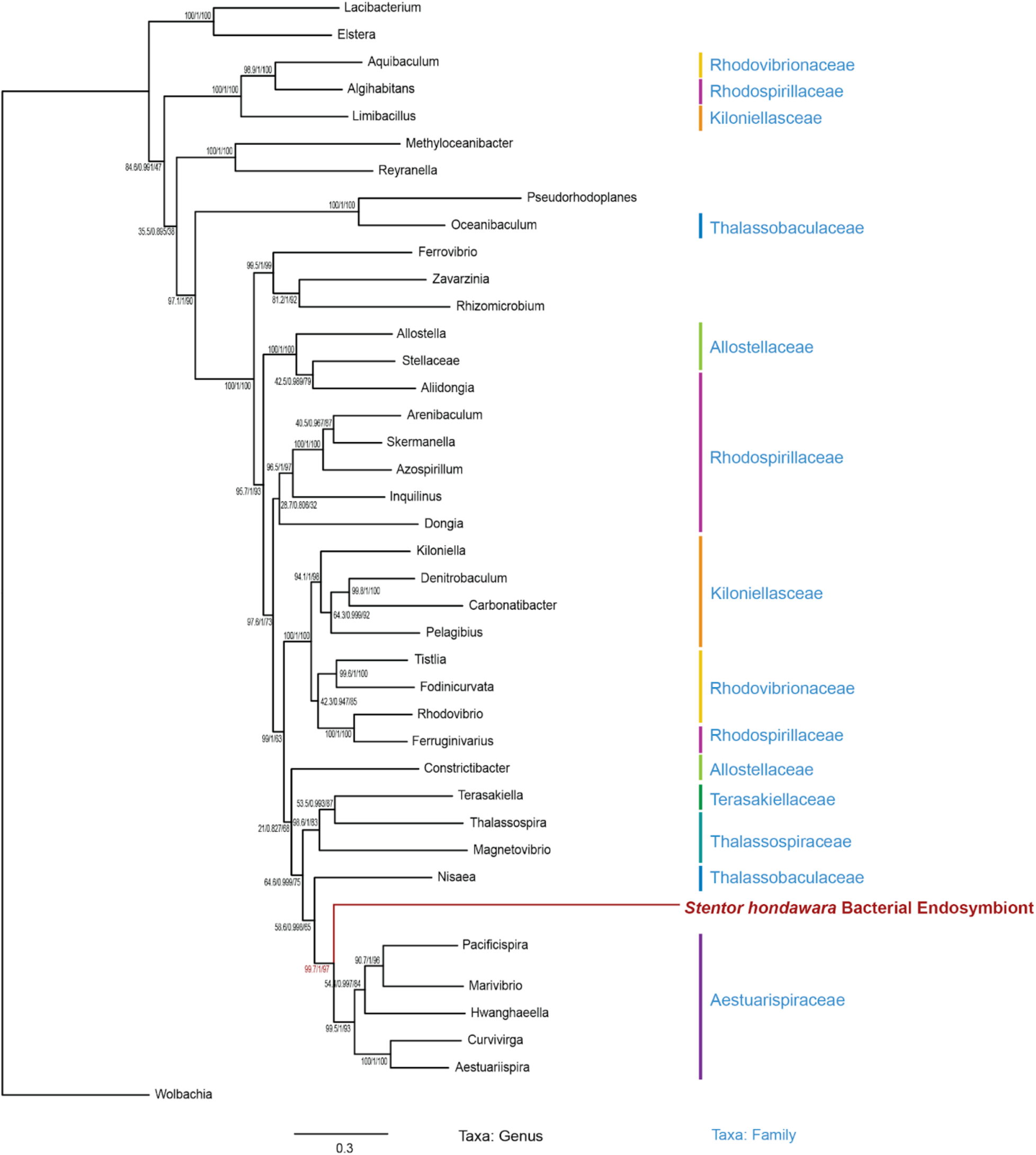
Phylogenetic tree of prospective endosymbiont with the order *Rhodospirillales* in the class *Alphaproteobacteria*. Phylogenetic tree assembled using IQtree based on maximum likelihood (ML) alignments of four genes, with branch lengths showing amino acid substitutions per site. A bacterium of the order *Rickettsiales* of the class *Alphaproteobacteria* serves as the outgroup. Scale bar indicates amino acid substitutions per site.

**Extended Data Table 1.** *Stentor hondawara-*specific enriched GO terms

**Extended Data Table 2.** Freshwater *Stentor* enriched GO terms

**Extended Data Table 3.** List of functionally-annotated genes from the mitochondrial genome of *Stentor hondawara*

**Extended Data Table 4.** List of functionally-annotated genes from the genome of a potential endosymbiotic bacteria in the *Stentor hondawara*

## Supplementary information

**Supplementary Data 1.** *Stentor hondawara* 18S SSU trimmed sequence

**Supplementary Data 2.** *Rhodospirillales* protein alignment sequences

**Supplementary Video S1.** Video of collection environment of *Stentor hondawara*

**Supplementary Video S2.** Video of active foraging behavior of *Stentor hondawara* in seawater

**Supplementary Video S3.** Video of contraction motions of *Stentor hondawara* in seawater

**Supplementary Video S4.** Video of active behavior and morphology of *Stentor hondawara* in seawater

**Supplementary Video S5.** 3D protein structure of *Stentor hondawara* ABCD4

**Supplementary Video S6.** 3D protein structure of *Stentor hondawara* CLCN2

**Supplementary Video S7.** 3D protein structure of *Stentor hondawara* Aquaporin

**Note:** The Supplementary Video files are listed only by title because of the *bioRxiv* guidelines’ allowable limit on total file size. All video files will be available in the peer-reviewed published version of the journal, which will be linked to the *bioRxiv* version accordingly.

